# Large scale prospective evaluation of co-folding across hundreds of Mac1-ligand complexes and three virtual screens

**DOI:** 10.64898/2025.12.25.696505

**Authors:** Jongbin Kim, Galen J. Correy, Brendan W. Hall, Moira M. Rachman, Olivier Mailhot, Takaya Togo, Ryan L. Gonciarz, Priyadarshini Jaishankar, R. Jeffrey Neitz, Eric R. Hantz, Yagmur U. Doruk, Maisie G. V. Stevens, Morgan E. Diolaiti, Rashad Reid, Saumya Gopalkrishnan, Nevan J. Krogan, Adam R. Renslo, Alan Ashworth, Brian K. Shoichet, James S. Fraser

## Abstract

Accurate prediction of ligand-bound protein complexes and ranking them by affinity are central problems in drug discovery. While deep learning co-folding methods can help address these challenges, their evaluation has been hampered by the difficulties in assessing independence from training data and insufficiently large test sets. Here we test the ability of co-folding methods to predict the structures of 551 ligands, 489 of which are of sufficient quality for evaluation, bound to the SARS-CoV-2 NSP3 macrodomain (Mac1) that were determined after the training cut-off dates. AlphaFold3 (AF3), Boltz-2, and Chai-1 each reproduced >50% of the Mac1 ligand poses to better than 2 Å RMSD of experiment. Despite the potential for co-folding to describe protein conformational changes that stabilize ligand binding, we did not find that common conformational rearrangements, including peptide flip and a large loop opening, were recapitulated by the co-folding prediction. For AF3 and Chai-1, ligand pose prediction confidence weakly, but significantly, tracked experimental potency, while DOCK3.7 energies were only weakly correlated. Boltz-2 affinity predictions showed the strongest correlation with measured potency and, after calibration, achieved lower mean absolute error than a baseline predictor. We next assessed whether co-folding scores could rescore docking hit-lists to distinguish true ligands from non-binders among hundreds of molecules prospectively experimentally tested against AmpC β-lactamase, the dopamine D4 and the σ_2_ receptors. AF3 ligand pose confidence values did not separate true ligands from high-scoring false-positives as effectively as docking scores or Boltz-2 affinity predictions did. Taken together, the modest, but independent correlations of docking score and co-folding confidence or affinity suggests that integrating physics-based and deep-learning and approaches may help with hit prioritization and subsequent optimization in structure-based ligand discovery.

## Introduction

Deep Learning methods such as AlphaFold2 (Jumper et al. 2021) and RosettaFold (Baek et al. 2021) revolutionized protein structure prediction. Key to their wide adoption has been their extensive testing, first in well-controlled prospective tests like CASP (Kryshtafovych et al. 2021; Ashizawa et al. 2025) and subsequently in many studies where the methods found new applications and new experimental validation (Scardino et al. 2023; Goverde et al. 2023). Notably, these methods can provide protein structural models for small molecule ligand discovery by docking/virtual screening, leveraging physics-inspired functions to score the resulting complexes (Seok and Tiwary 2025; Brocidiacono et al. 2025; Huang et al. 2006; Michino et al. 2025; Xie et al. 2025). Retrospective analyses of AlphaFold2 models suggested that the accuracy of the predicted ligand binding poses was lower than experimental structures (Karelina et al. 2023; Holcomb et al. 2023). However, prospective use of AlphaFold2 models and experimental structures (determined after the training cutoff) to identify new ligands revealed that both approaches yielded similar quality and quantity of ligands (Lyu et al. 2024; Díaz-Holguín et al. 2024). These results indicate that AF2 models may sample alternative low energy conformations that are compatible with distinct ligands from the experimentally resolved conformations, reconciling the poor retrospective recall with the prospective successes.

More recently, the advent of co-folding methods like AlphaFold3 (AF3) (Abramson et al. 2024), Chai (Boitreaud et al. 2024), and Boltz (Wohlwend et al. 2025) that leverage diffusion-based approaches for multi-component (e.g. protein and small molecule) complex predictions, have promised to extend the impact of deep learning methods from protein structure prediction to small molecule ligand-bound protein complexes (He et al. 2025). The advantages co-folding may have over traditional docking include its abilities to explore conformational variability of the protein on-the-fly and to predict ligand affinity, reportedly as well as or better than a gold-standard physics-based method, free energy perturbation (FEP) (Passaro et al. 2025). However, multiple studies have highlighted how co-folding success can track with structural similarity to training structures, consistent with substantial structural memorization (Xu et al. 2025; Škrinjar et al. 2025). An innovative approach introduced adversarial mutations to fill binding sites to show that co-folding models will still produce high confidence poses even when interactions are physically impossible (Masters et al. 2025). These results highlight pervasive memorization and hallucination in current co-folding methods and suggest that alternative computational approaches are needed to quantify the physical basis of the protein-ligand interactions.

Still, co-folding has potential for synergizing with many aspects of structure-based design and more traditional computational chemistry approaches. Two attractive applications are the prediction of ligand-bound complexes, around which structure-based ligand design is articulated (Erlanson et al. 2025); (Backus et al. 2016), and the re-scoring of large library docking hitlists. Co-folding is orders of magnitude slower than most docking programs, which can address libraries of hundreds-of-millions to billions of compounds on CPU clusters. The method nevertheless is fast enough to re-rank virtual screening “hit-lists” and may be better at categorizing certain types of ligands (e.g. those that stabilize alternative conformations) than the simpler physics-based methods. Moreover, the errors and biases from co-folding affinity estimation methods (Passaro et al. 2025) may be independent from those in docking, suggesting that each method may have different strengths in distinguishing likely from unlikely ligands. However, partly because of the lack of widely adopted community prospective tests, akin to CASP (Gathiaka et al. 2016) and partly because of the challenges of developing, testing, and potentially optimizing new chemical matter for ligand activity studies (Ackloo et al. 2022), co-folding methods have not yet been subject to the rigorous, large-scale testing that helped establish the utility, domain of applicability, and ultimately optimization of the early protein structure-prediction methods.

It therefore seemed interesting to test co-folding predictions of the co-complexed structures and affinities of ligands in a drug-discovery series from outside of their training sets, and to test their ability to distinguish likely from unlikely ligands from large library virtual screens (Cabeza de Vaca et al. 2025). Here, we evaluated the performance of three widely-used co-folding methods, AF3, Chai-1, and Boltz-2 against two benchmarks that speak to domain utility and are largely outside the benchmarks against which they trained. First, we tested the ability of the co-folding methods to recapitulate the crystallographic structures of 551 different ligands bound to the SARS-CoV-2 Macrodomain (Mac1), an emerging antiviral target (Suryawanshi et al. 2025). This dataset represents the scale and depth typical of the early phase of industrial structure-based drug design campaigns. Second, we investigated the ability of the co-folding methods to distinguish true-from false-positives in hit lists from recent large library, prospective docking against three well-behaved targets: AmpC β-lactamase, the σ2 receptor, and the dopamine D4 receptor (Lyu et al. 2019; Alon et al. 2021; Liu et al. 2025). In the campaigns against these targets, between 500 and 1500 molecules were prospectively tested across a range of docking ranks identifying ligands with previously unexplored chemotypes as well as finding many docking false positives. Contemporaneous work has highlighted the importance of using AF3 in the same experimental datasets from ultra-large library docking campaigns (Menon et al. 2025). Our analyses reveal the strengths and limitations of different co-folding methods versus traditional docking and point to potential synergies that can improve probe and lead discovery pipelines.

## Results

### A set of 551 previously unreported Mac1 ligands complexes

As part of a drug discovery effort (Suryawanshi et al. 2025; Jaishankar et al. 2025; Correy et al. 2025; Flowers et al. 2025; Gahbauer et al. 2023; Schuller et al. 2021), we tested thousands of novel small molecules as inhibitors of Mac1, a new target for SARS-CoV2 and other viruses. This led to potent inhibitors that are active *in vivo* as anti-virals (Suryawanshi et al. 2025; Jaishankar et al. 2025). As part of this work, we determined x-ray crystal structures for ∼1000 small molecules bound to Mac1, typically at close to 1 Å resolution, but only a limited number of these were disclosed in the papers describing the more advanced candidates. Moreover, only up to 326 of our molecules, largely consisting of initial fragments and early hit-to-lead material, were available in public databases prior to the training cutoffs for the cofolding methods (e.g. 2021-09-30 for AF3, 2023-06-30 for Boltz-2). Apart from their role in driving inhibitor optimization for Mac1, we thought that these structures might be a resource for evaluating computational approaches against a set that is representative of the scale and depth often seen in industrial structure-based drug design campaigns.

From our internal dataset, we identified 551 ligands where we determined X-ray structures that have not been disclosed previously (**Fig. 1**a, **Supplementary File 1**). These molecules all bind in the active site where they compete with the substrate, ADP-ribosylated (ADP-r) peptides. There are several stereotyped interactions across this set, which have been described in previous fragment and lead optimization efforts (Gahbauer et al. 2023; Schuller et al. 2021; Correy et al. 2025), including: interactions with backbone NH groups of Phe156 and Asp157 (the oxyanion site), aromatic packing against the side chain of Phe156 (mimicking the adenosine ring of ADP-r), and hydrogen bonds to Asp22 and Ile23 (the upper active site) (**Fig. 1**a). Because these ligands were synthesized during an active campaign and many did not meet evolving thresholds for single point % inhibition, only 202 have fully measured dose response IC_50_ estimations by a HTRF-based peptide displacement assay. For these compounds, the measured potencies range from sub-micromolar to >500 uM (**Fig. 1**b). This distribution is skewed towards weaker affinities as more potent molecules were progressed and disclosed in papers describing *in vivo*-active lead molecules (Suryawanshi et al. 2025; Jaishankar et al. 2025).

**Figure 1:**
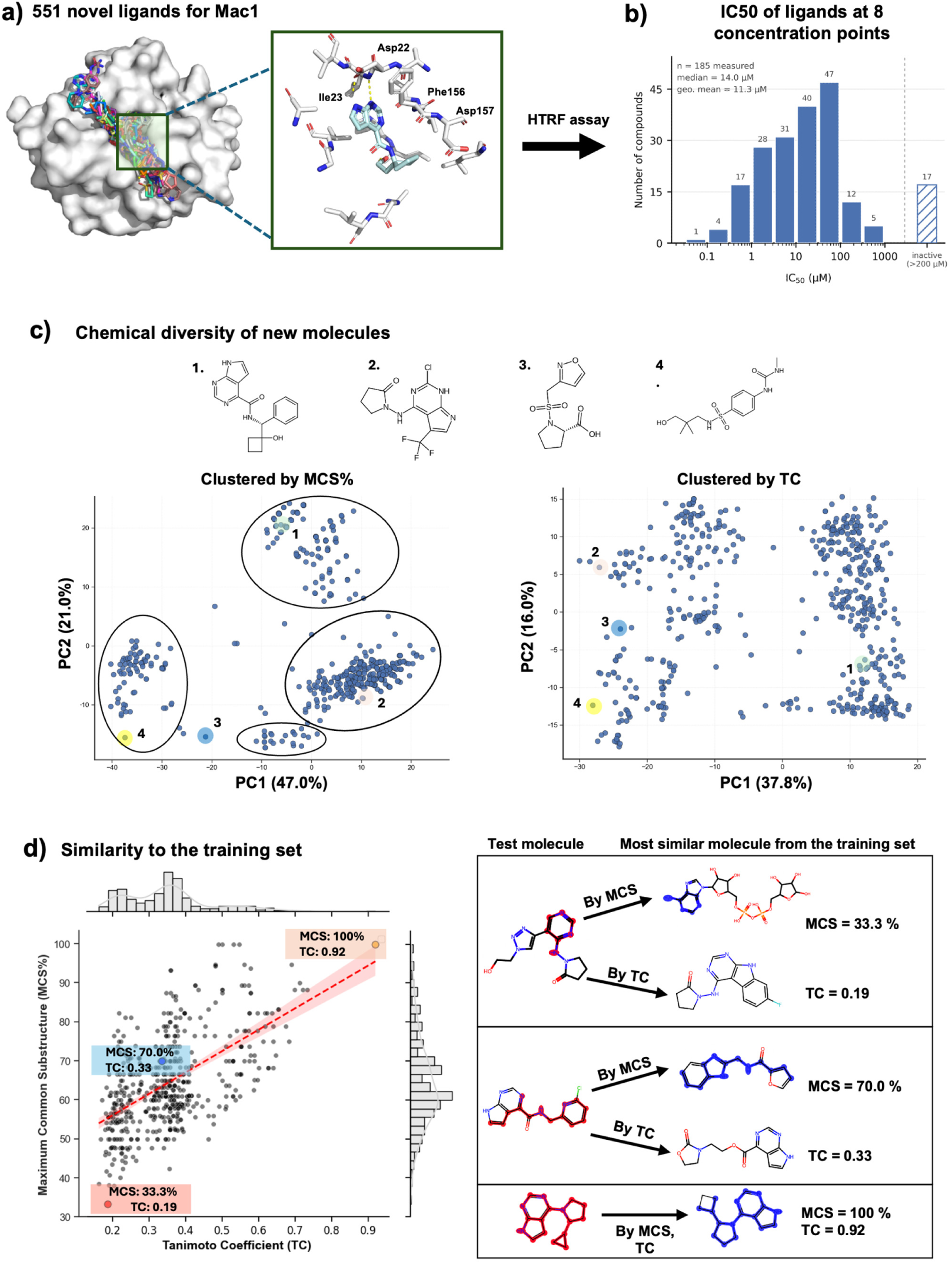
Structure diversity of 551 previously unreported ligands co-folded with the SARS-CoV-2 Mac1 (NSP3 macrodomain 1). **a)** Global superposition of all 551 ligands within the Mac1 binding pocket highlighting residues involved in key interactions. **b)** The affinity distribution of 202 molecules determined by HTRF (drug-response curves at 8 concentration points). **c)** Principal Component Analysis projection of compounds represented by their full pairwise similarity vectors (by ECFP-4 Tc and MCS%). **d)** The highest MCS% vs. ECFP-4 Tc values in the training set for each Mac1 compound. At right are examples of the low similarity, scaffold hop, and high similarity distributions, respectively.

The diversity of the 551 ligands is important for understanding the performance of the cofolding methods. By topological similarity, using ECFP4-based pairwise Tanimoto coefficients (Tc), the ligands were diverse, with pairwise Tc clustering between 0.2 to 0.4 (**Figure 1—figure supplement 1a**, **Supplementary File 2**). For context, a scaffold-hop for the ECFP4-based fingerprint is thought to occur at Tc values below 0.35 (Cramer et al. 2004; Muchmore et al. 2008), while random similarity occurs around Tc values of 0.25 (Hert et al. 2008). As the Tc metric has well-known liabilities that can mask what a chemist would see as similarity, we also calculated pairwise Maximum Common Substructures (MCS) and found a distribution that centers around 44.8% (**Figure 1—figure supplement 1b, Supplementary File 2**). Clustering based on pairwise MCS > 35% revealed only four distinct cluster heads and a limited number of outliers (**Fig. 1**c, **Supplementary File 2**, **Figure 1—figure supplement 2**). To quantify the potential for model performance to be driven by memorization, for each ligand in our set, we retrieved the most similar ligand that was deposited prior to the training cutoff date by both MCS and Tc (**Fig. 1**d). Here, the Tcs have a trimodal distribution with a small number having Tc > 0.4 (indicating relatively high similarity for this fingerprint), and two larger distributions centered around 0.35 (indicating scaffold hops) and 0.2 (indicating low similarity to any ligand). The MCS similarities spanned a broad range centered around ∼60%, consistent with the majority of the dataset representing a small number of conserved scaffolds bearing diverse peripheral substitutions. We could also confirm the chemical diversity of the Mac1 compounds relative to the broader chemical space represented by all ligands deposited in the PDB by both Tc and MCS% (**Figure 1—figure supplement 1c-d**).

### Co-folding can accurately reproduce poses of ligands dissimilar to those trained

Next we assessed how well co-folding methods could predict the structures of the 551 complexes. Three co-folding methods (AlphaFold3, Chai-1, and Boltz-2) were given the SMILES strings of each ligand and the sequence of the enzyme as inputs. In parallel, we performed classic docking with DOCK3.7 using the SMILES of the ligand and the three dimensional structure of the enzyme bound to an early inhibitor (PDB: 5SQW, (Gahbauer et al. 2023) that is roughly contemporaneous with the ligands in this datasets. However, 68 structures had weak density for part of the molecule, and were not used as reference structures for comparing co-folded poses. To assess whether ligands were correctly predicted in the right area of the active site (Nittinger et al. 2025), we used a threshold of ligand center of mass < 2.5 Å after alignment of 489 protein structures: 449 AlphaFold3 poses (92%), 440Chai-1 poses (90%) and 444Boltz-2 poses (93%). With DOCK3.7, only 80% (392 out of 489 poses) passed this cutoff. In addition, roughly 1.5% of co-folded or docked molecules did not pass the RDkitbased sanitization pipeline necessary for RMSD comparisons. Therefore, the total number of poses compared for prospective prediction differed slightly among all co-folding and docking methods (**Supplementary File 3**).

Across the co-folding methods, AF3 and Chai-1 performed qualitatively similar with a roughly unimodal distribution centered around ∼1 Å (the mode) RMSD values (**Fig. 2**a). Using the field standard, 2 Å cutoff for correct predictions, AF3 and Chai-1 predicted roughly70% of the ligands correctly. Boltz-2 performed less well, with more structures shifting to the 2 Å distribution, and an overall predictive accuracy of 53%. This result was surprising because the later training cut-off date for Boltz-2 meant that it could potentially benefit from additional Mac1 ligand-bound structures. The distribution for docking was more bimodal centered around an RMSD of 2 Å and 5 Å, yielding a predictive accuracy of 40%. At least for this set, the co-folding models outperformed DOCK3.7 in accurately predicting the poses of bound ligands.

**Figure 2:**
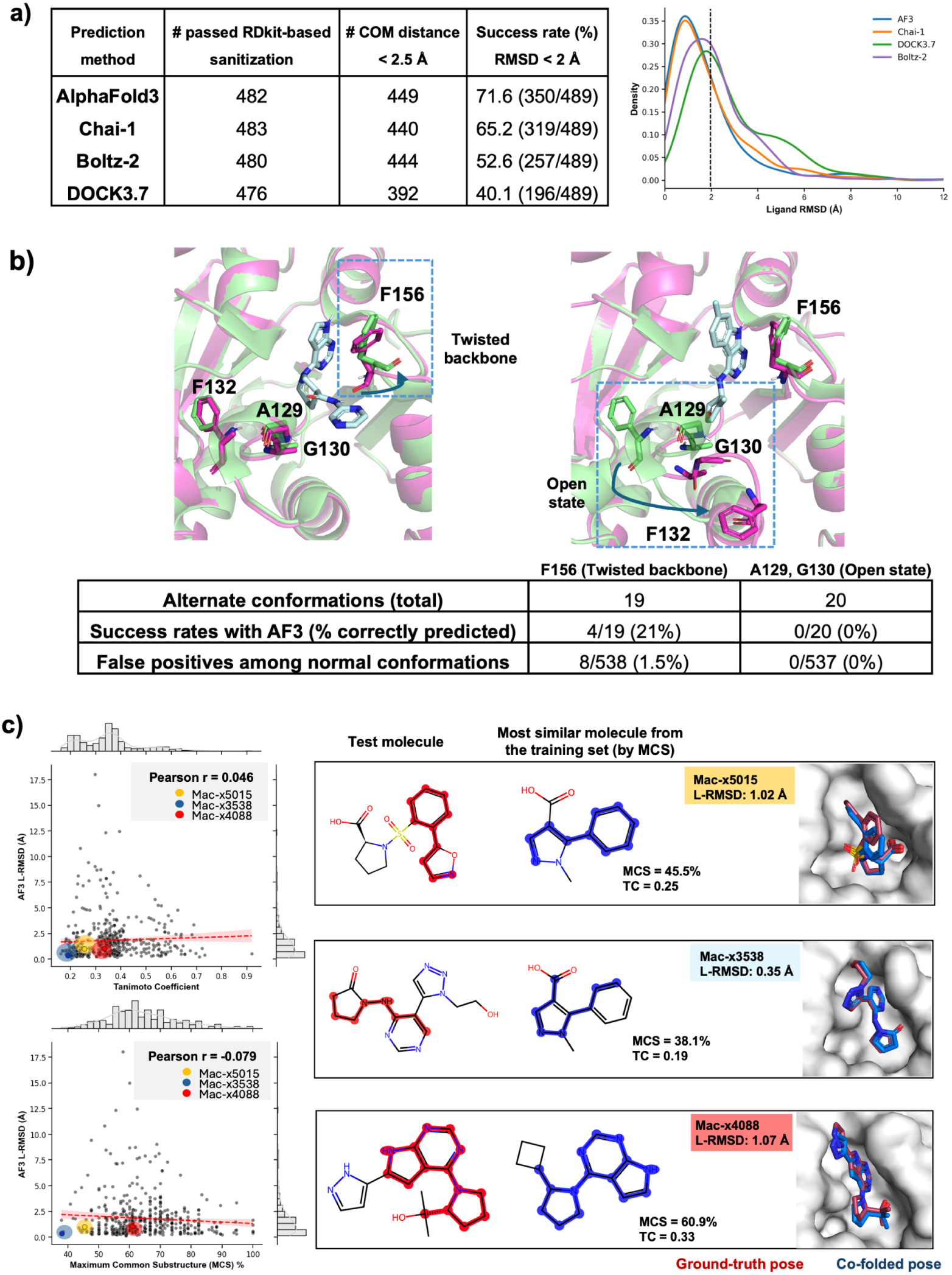
Co-folding outperforms DOCK3.7 in predicting the ligand pose for 489 ligands against Mac1 binding site. **a)** Histogram distributions of ligand heavy-atom RMSD (L-RMSD) between predicted and crystallographic poses obtained with AF3, Chai-1, Boltz-2 and DOCK3.7, and the success rate defined as RMSD < 2 Å, b) Demonstration of alternate conformations of Mac1 binding pocket (twisted backbone and open state) and ability of AF3 to capture these conformational changes (residues in green are from PDB ID: 5SQW, and residues in pink have altered conformations), c) AF3 L-RMSD pose recovery is compared with TC ECFP4 and MCS% and accurate poses for test molecules with poor similarity to the closest molecule in the training set.

Next we investigated the potential for co-folding methods to identify alternative conformations of the protein that are stabilized by ligand binding. This application is a potentially large differentiator relative to virtual screening, which generally uses a fixed structure. Although protein alignment gave generally low global C-alpha RMSDs (range 0.1 Å – 0.4 Å; **Supplementary File 3**), rare local and high magnitude conformational changes observed experimentally were not captured by co-folding methods. Previously published structures indicated two characteristic states with significant conformational changes: a twisted state, characterized by a peptide backbone flip around Phe156, and an open state, characterized by a ∼8 Å loop rearrangement including Gly130 and Ala129 (**Fig. 2**b, **Figure 2—figure supplement 1**, **Supplementary File 4**). The open state has also been observed in the human MacroD1 enzyme (PDB 2X47, (Chen et al. 2011) and PDB 6LH4, (Yang et al. 2020)), suggesting it might be a common feature across macrodomains. We observed that only 4/19 twisted states and 0/20 open states were correctly predicted by AF3. Interestingly, AF3 predicted 8 false positive twisted states, where the ligand was experimentally observed in the normal ground state. This result suggests that there is only a limited ability to predict conformational adaptations, which admittedly may be, at least in part, stabilized by crystal contacts. We next asked how often the predictions recapitulated hydrogen bonding interactions in the Mac1 active site (**Figure 2—figure supplement 1a**). Among the dominant interactions, we found a low false positive rate for hydrogen bonds with D22 and I23 at the top of the active site, and mispredictions in both directions for hydrogen bonds with F156 and D157 at the oxyanion site (**Figure 2—figure supplement 1b**). Thus, despite failing to predict large conformational changes and aspects of the detailed hydrogen bonding interactions, AF3 still correctly placed these ligands with low RMSDs relative to the crystal structure pose (Errington et al. 2025). This result suggests that co-folding reliably recapitulates dominant ligand-binding interactions even in the absence of accurate protein conformational modeling, providing further support to the idea that they are learning specific interaction patterns rather than a deeper physics-based representation (Masters et al. 2025).Finally, we asked how much of co-folding success might be explained by memorization of Mac1 ligands deposited prior to the training date (**Fig. 1**d). We measured the correlation between AF3 ligand RMSD, and either the Tc (ECFP4) or %MCS of the most similar ligand within the training set, finding no meaningful signal (**Fig. 2**c). Examples illustrated have low similarity in both metrics, and mac-x5015, mac-x3538 are not from the commonly shared purine scaffold. Nonetheless, even for these cases, AF3 co-folded poses are recovered to within 1 Å accuracy. The weak correlations between RMSD and both similarity metrics illustrate the ability to predict binding modes for ligands dissimilar from the training set.

### Model confidence and energy scores reflect pose reproduction and affinities

A central question for any co-folding or docking method is whether its internal confidence or energy scores track with experimental pose accuracy and affinity. We used the structure confidence scores reported by AF3, the per atom Predicted Local Distance Difference Test (pLDDT) scores for ligand atoms, and for Chai-1 the interface predictive template modeling (ipTM) score. While these metrics are intended to estimate the model confidence in accuracy, they are not explicitly designed to predict affinity. In contrast, the Boltz-2 affinity module predicts IC_50_ (pIC_50_) which should correlate with affinity, whilst the DOCK3.7 score, though in kcal/mol, is not expected to correlate strongly with experimental affinity (Shoichet 2004; Irwin and Shoichet 2016; Tirado-Rives and Jorgensen 2006). Boltz-2 affinity score and DOCK energies do not explicitly estimate the accuracy (RMSD) of the prediction.

We first generated Receiver Operating Characteristic (ROC) curves to measure the retrieval of True Positives (good score, RMSD < 2Å) versus False Positives (good score, RMSD > 2Å) (**Fig. 3**a). We observed the best performance for AF3 L-pLDDT (AUC=75.6) and Chai-1 ipTM (AUC=74.6). This indicates that the internal model confidence has predictive power and, indeed, we observed that these scores are correlated with RMSD (**Figure 3—figure supplement 1**). Perhaps not surprisingly, the performance of the DOCK3.7 energy (AUC=67.1) and Boltz-2 pIC_50_ (AUC=53.8) scores, which are designed to predict affinity, was poorer than those designed to predict accuracy. The DOCK3.7 scores correlated with Mac1 ligand RMSD with a Pearson *R* of 0.33(p < 0.001), which is in line with expectations from historical analyses of docking studies (Wierbowski et al. 2020). However, Boltz-2 affinity scores did not correlate with the predicted pose (Pearson r = −0.001, p > 0.5), which likely results from the fact that its internal structure prediction confidence module is separated from the affinity module (Passaro et al. 2025). Indeed, using the Boltz-2 structure prediction confidence metric (Ligand ipTM), yields an AUC of 75.2, comparable to AF3 and Chai-1. Considering its early enrichment performance, Boltz-2 Ligand ipTM was the strongest predictor of pose accuracy based on normalized logAUC (27.1% above random, **Fig. 3**a). In contrast, although Boltz-2 pIC50 showed poor overall discrimination, it overestimated its ability to enrich true positive poses at low false positive rates, despite having a weak early enrichment behavior. Since confidence scores correlated with RMSD, we also investigated how each method was able to predict poses and scores depending on similarity of molecules to the training set (**Figure 3—figure supplement 2**). As before (**Fig. 2**c), the performance of co-folding and docking scores did not show a strong dependence on novelty relative to the training set. This result suggests that co-folding can accurately reproduce poses of ligands dissimilar to those on which it trained, and that the model internal confidence scores reflect how well poses are reproduced.

**Figure 3:**
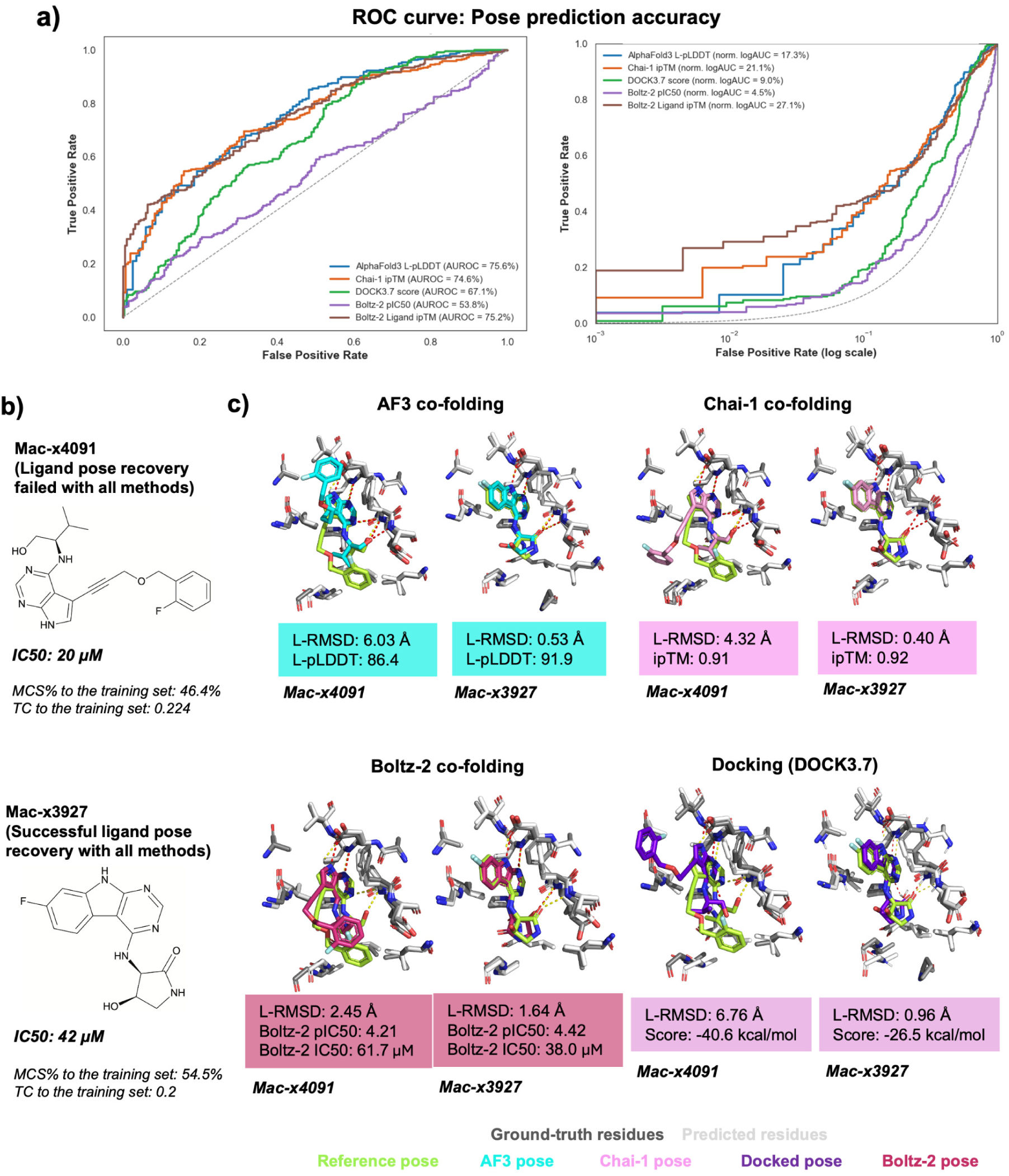
Complementarity between co-folding confidence metrics and ligand-pose accuracy. **a)** Receiver Operating Characteristic (ROC) curves for affinity/energy and confidence scores in AlphaFold3, Chai-1 and Boltz-2, with the success rate of binding pose prediction being defined as L-RMSD < 2 Åand both AUC and logAUC are shown in percentages. b) Chemical structure of a compound that was challenging to model with all docking, co-folding methods (mac-x4091), and one that was well-modelled with all methods (mac-x3927), c) Overlay of co-folded (AlphaFold3, Chai-1, Boltz-2) and docked ligand poses on ground-truth, for hits with similar IC_50_, but different structures. Hydrogen bonds with co-folded, docked ligands are shown in red, and those with crystal ligands are shown in yellow.

Representative examples of correct and incorrect pose predictions across co-folding and docking methods are illustrated with mac-x3927 and mac-x4091, respectively (**Fig. 3**b,c). Both compounds exhibited comparable experimental IC_50_ values (20 µM for mac-x4091 and 42 µM for mac-x3927) and similarity to molecules in the training set (**Fig. 3**b). The molecule consistently predicted correctly (mac-x3927) was a small, rigid fragment comprising a purine scaffold rigidly linked to a fluorobenzene moiety, favoring a well-defined binding mode. In contrast, the compound mispredicted by all methods (mac-x4091) was larger and more conformationally flexible, with an alkyne linker prone to flipping, potentially altering the orientation of the fluorobenzyl ether substituent. Although this group packs against a crystal contact, the high predictive confidence/affinity scores across co-folding methods (AF3 L-pLDDT = 86.4, Chai-1 ipTM = 0.91) are likely explained by the adenosine mimic purine group where predictions recapitulate key interactions observed in the experimental structure.

To understand if prediction errors between methods are correlated, we directly compared the agreement between AF3 to Chai-1, Boltz-2 or DOCK3.7 predictions with the agreement between AF3 and experimental ground truth (**Figure 3—figure supplement 3**). We observed that while there are a small number of structures where AF3 predicts much better than other methods (high AF3-other prediction RMSD, low AF3-experiment RMSD), most large errors are either missed in a similar way (low AF3-other prediction RMSD, but high AF3-experiment RMSD) or are both extreme mispredictions (high AF3-other prediction RMSD and AF3-experiment RMSD). Consistent with this result, in comparing RMSD relative to experimental structures for all methods (**Figure 3—figure supplement 4**), we observed that Chai-1 and AF3 exhibited the strongest agreement (r = 0.72), whereas poses from DOCK and Boltz-2 diverged more (r = 0.45 for AF3 vs. DOCK, and r = 0.52 for AF3 vs. Boltz-2). Collectively, these results suggest that the co-folding methods (AF3, Chai-1) may be complementary to more physics-based approaches like DOCK3.7 as the errors are less correlated than observed for the different cofolding methods.

Second, we asked how well these scores correlate with experimentally measured potency (**Fig. 4**a). The correlation was best for the Boltz-2 pIC_50_ (r = 0.6), significantly outperforming DOCK when tested with Friedman and Conover tests with Bonferroni corrections (p = 0.0057). Prior to any linear correction, the mean absolute error (MAE) in affinity prediction was ∼1.6 kcals (1.19 pIC units), with most affinities predicted as too potent. Empirically correcting with a linear fit reduced this error to ∼0.7 kcals (0.54 pIC units). To put these results in context, simply predicting the mean pIC50 from the dataset for every compound would yield a baseline MAE of 0.93 kcal because of the relatively limited dynamic range of the dataset. The DOCK3.7 energy correlated more weakly with potency (r = −0.225), consistent with previous studies that have observed only weak correlations with affinity from virtual screening energies (Kinnings et al. 2011; Jones et al. 2021). The AF3 co-folding confidence metrics correlated slightly better than docking with potency (r = 0.314, **Fig. 4**a), but the post-hoc Conover tests with Bonferroni corrections showed the difference was not statistically meaningful. Despite this, a correlation between pose accuracy and potency was more relevant with AF3 (r = −0.233, **Fig. 4**b), as more accurate poses tend to be more confidently predicted (**Figure 3— figure supplement 1**). In contrast, there was no correlation between docking accuracy and potency (**Fig. 4**b). Boltz pIC_50_ had the highest correlation between pose accuracy and affinity (r = −0.279). This result is interesting because in Boltz-2 affinity prediction and confidence are decoupled architecturally and have little correlation with each other across the Mac1 dataset (Figure 3—figure supplement 1).

**Figure 4:**
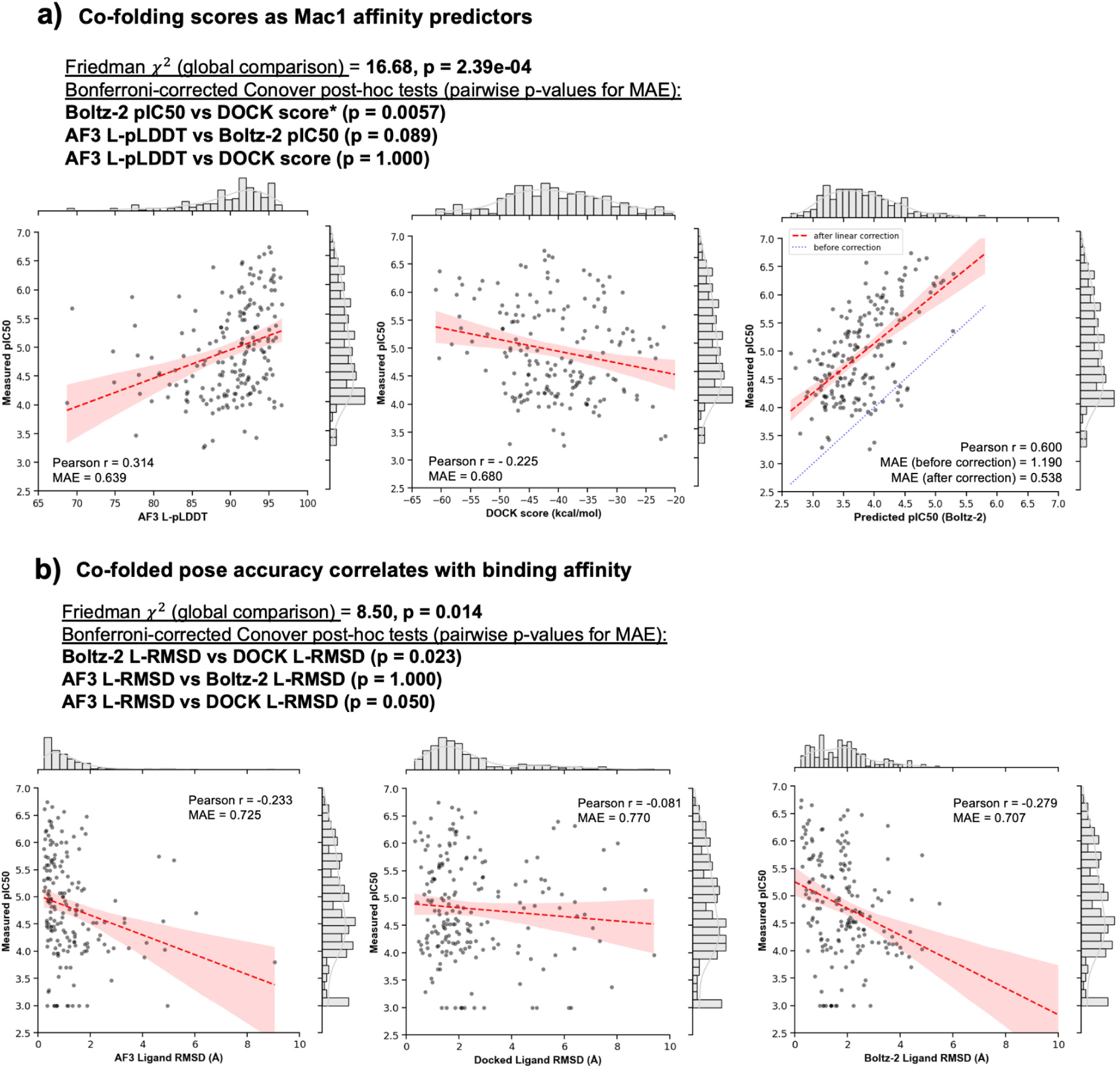
Co-folding scores and pose accuracy correlate with Mac1 binding affinity for 202 newly synthesized Mac1 ligands. **a)** AF3 ligand pLDDT (L-pLDDT) and Boltz-2 predicted pIC50 correlate with experimental pIC50 more strongly than DOCK energy scores, b) Lower co-folded ligand RMSDs associate with higher affinity, while docked pose RMSDs do not. Pearson’s Correlation Coefficients and Mean Absolute Errors (MAE) are shown. The baseline MAE was 0.68 pIC (or 0.93 kcals), measured against a fixed average pIC50 (pIC50 = 4.95). Friedman tests with Conover post-hoc comparisons and Bonferroni corrections are used to compare MAE between methods.

Collectively, these results suggest that the categorization abilities of the co-folding confidence metrics, and their correlation with affinity, could mark an important advance. Still, the Mac1 set, as large as it is, represents only a single system with ligands that share a limited set of pharmacophores. We wanted to broaden testing to more targets and more diverse ligands, such as those encountered in virtual screening hit lists with their focus on new chemotype discovery.

### Co-folding struggles to reliably re-rank Docking Hit-lists

Encouraged by the ability of co-folding to reproduce ligand structures and affinities, we were interested to see if the methods could also improve hit identification from large library docking screens. In these screens, hundreds of millions to billions of molecules are docked against a target and scored for complementarity to a binding site. High ranking molecules are then synthesized and tested for binding. While large library docking has successfully discovered potent novel chemotypes in recent studies (Lyu et al. 2019; Gorgulla et al. 2020; Stein et al. 2020; Sadybekov et al. 2022; Fink et al. 2022; Singh et al. 2023; Liu et al. 2025), the method is plagued by false positives (molecules that rank well but are inactive). We reasoned that if co-folding can predict the structures and activities of true ligands confidently, perhaps it could distinguish docking true binders from false positives.

To test this idea, we analyzed experimentally tested compounds from three large and heavily tested docking campaigns (Supplementary File 5): 506 experimentally-tested compounds from a 490-million molecule screen against the σ_2_ receptor (Alon et al. 2021), 1,293 tested compounds from a 1.7-billion molecule screen against AmpC β-lactamase (Liu et al. 2025), and 541 tested compounds from a 138-million molecule screen against dopamine D4 receptors (Lyu et al. 2019) with AF3 (the best performing model from the Mac1 set), DOCK scoring function, and the Boltz-2 pIC_50_ score. These test sets differ from the Mac1 benchmark in three ways. First, they are a mix of molecules that would turn out to be true binders (247 in AmpC, 201 in σ_2_ and 205 in Dopamine D4) and high-ranking docking false positives as judged by measured potencies (worse than 400 μM in AmpC, worse than 2.0 μM against the σ_2_ receptor, and worse than 7.8 μM against the dopamine D4 receptor). For these molecules, the challenge is not only predicting the bound structures of true ligands, but distinguishing true ligands from decoy molecules that by many biophysical criteria fit the binding site well. Second, the docking molecules are more diverse than the Mac1 set (**Figure 1—figure supplement 2**), which largely falls into four substructure clusters representing conserved pharmacophores. The docking sets were selected, in the original studies, for their topological diversity, and their dissimilarity to previously known ligands. For each target, the ligands group into more than 10-times the number of clusters than the Mac1 dataset, consistent with a greater diversity. Indeed, the average pairwise ECFP4 Tc values within the three sets vary from 0.12 to 0.15, all well-below scaffold-hop cut-offs for this fingerprint (**Figure 1—figure supplement 2**, **Supplementary File 2**). Finally, the sets were all selected for novelty and there was little similarity between the ligands in the hit-lists and known ligands, which ranged between 0.11 and 0.19 by Tc, and between 33 to 42 by MCS%. These ranges were lower than our prior Mac1 dataset (where Tc is distributed around 0.36, and MCS% is distributed around 60.1%), and the lack of similarity with the training set for target systems with hit-lists could be explained by having relatively fewer ligand structures to compare to in the PDB (**Figure 4—figure supplement 1**, **Supplementary File 6**). These conditions, while representing new and difficult tests for co-folding, are the common case for structure-based ligand discovery campaigns. For validation, there are more measured IC_50_ values, but they are admittedly not as deeply supported by structural information as the Mac1 set.

We observed the strongest discrimination between active and inactive molecules in the σ_2_ dataset, which contains 201 confirmed actives (apparent *K_i_*values between 2.5 nM and 1.5 µM) and 305 inactive false positives (**Fig. 5**a). DOCK3.7 scores well-separated binders from non-binders (median –55.0 vs –42.5 kcal mol^-1^; AUC = 78.8), though the correlation between docking score and *pK_i_* for true ligands was relatively weak (r = –0.211). AF3 L-pLDDT showed a poorer ability to discriminate between true ligands and false positives (median 75.9 vs 74.1, AUC = 56.1). Intriguingly, despite the worse discriminatory ability, the AF3 correlation with binders *pK_i_* was not much worse than docking’s (r = 0.183). Against the σ_2_ set, Boltz-2 pIC_50_ performed better still, with high discriminatory power (median 6.70 vs 5.38, AUC = 83.8) and the strongest correlation of affinity predictions against true positives (r = 0.486). Among active molecules, Boltz-2 pIC_50_ had a MAE of only 0.7 kcal and a linear calibration reduced that error to 0.54 kcal (**Figure 5—figure supplement 1**). This absolute estimation of IC_50_ was superior to its performance by ∼0.5 kcal on the Mac1 dataset; however, it is restricted only to true positives and the performance could be much worse if the false positives were accounted for in the comparison. Since inactive predictions were in the same pIC_50_ range as active molecules, as an optimistic limit of the performance that includes inactive molecules, we assigned an IC_50_ of 4 µM, which is 2X our threshold for declaring a non–binder, to all false positives and calculated the MAE. With this assumption, the MAE is 0.92 kcal (0.56 kcal after linear calibration). A less generous assumption would be to randomly assign an IC_50_ between the *pK_i_* threshold (2 µM) and *pK_i_* = 1 (100 mM). Under this assumption, the MAE is 1.66 kcal (1.55 kcal after linear calibration) (**Figure 5—figure supplement 2**), which is in line with previously published estimates of the accuracy of this method (Passaro et al. 2025) and only a slight improvement over the MAE (1.84 kcal) produced by predicting the dataset average affinity for each datapoint.

**Figure 5:**
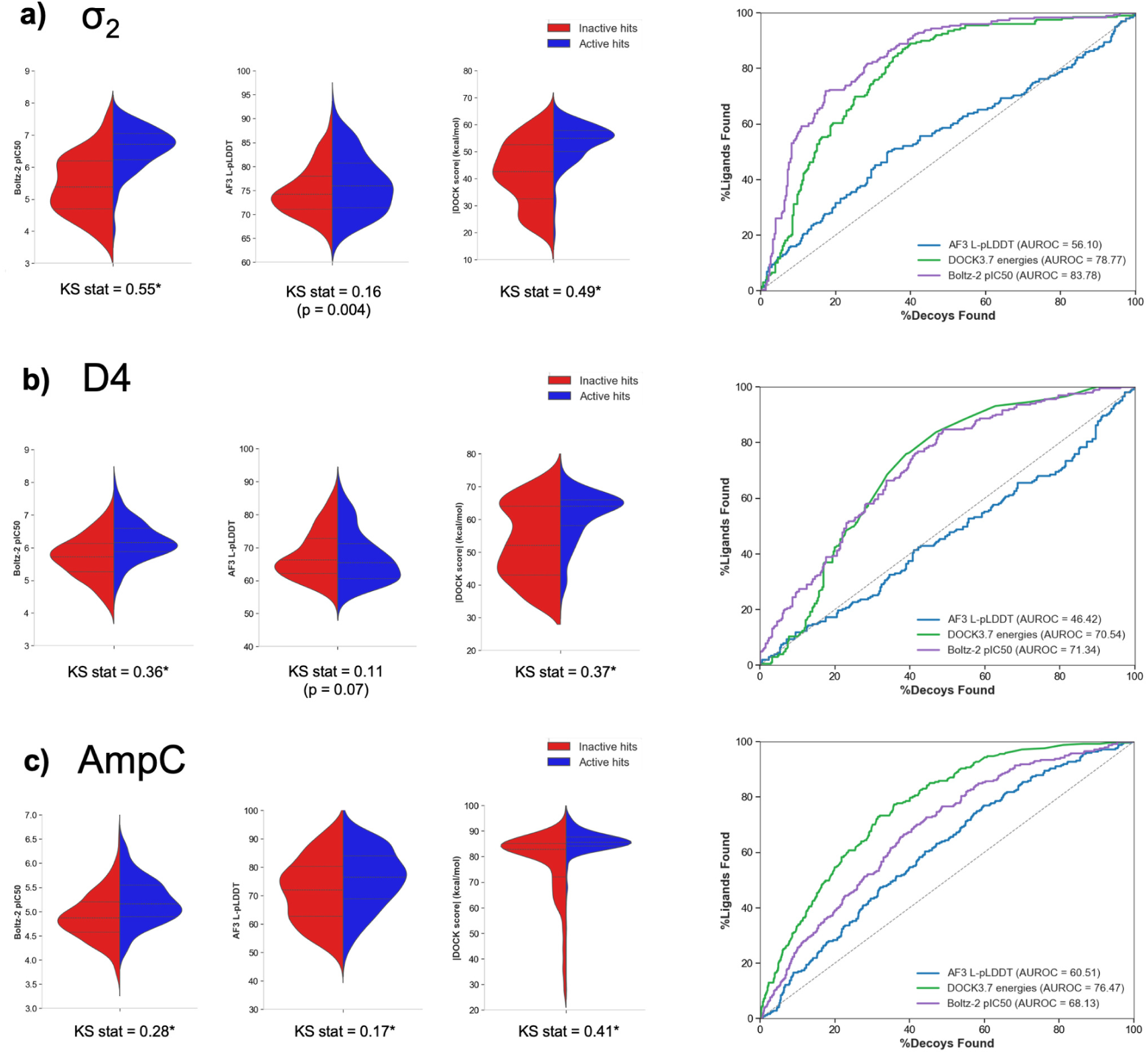
Classification of known active hits from docked false positives from an initially-screened docked molecules. The leftmost panel shows affinity scores from Boltz-2 (Boltz-2 pIC_50_), the middle panel for AF3 L-pLDDT and the right panel for absolute value of DOCK energies (kcal/mol) from DOCK3.7. Docked hit lists from a) σ_2_, b) Dopamine D4, c) AmpC are grouped into known hits and non-hits, and score distributions are shown in violin plots. Komolgorov-Smirnov test and its KS-statistic (KS stat) is used to indicate meaningful separation of two distribution curves. Asterisks indicate the p-value of KS statistic < 0.001. ROC curves show an ability to distinguish known actives from the docked list with different scoring methods.

For AmpC (**Fig. 5**b), a total of 1,293 high-ranking molecules were experimentally tested for inhibition (Liu et al. 2025). Among these were 247 molecules that turned out to be true inhibitors. We observed that docking (AUC = 76.5, median −85.6 vs −82.8 kcal/mol) was much stronger than Boltz-2 (AUC = 68.1, median 5.16 vs 4.87) or AF3 (AUC = 60.5, median 76.6 vs 72.0) in discriminating actives from inactives. The correlation of metrics to *pK_i_*among actives was relatively weak for DOCK3.7, AF3 and Boltz-2 in this case. The Boltz-2 pIC_50_ values for AmpC were particularly high in absolute error 1.81 kcal; however, linear correction reduced this error to only 0.42 kcal. Including inactive molecules degraded performance to 2.33 kcal (0.32 kcal with linear correction) under the optimistic and 3.29 kcal (1.04 kcal with linear correction) under the random IC_50_ assumptions (**Figure 5—figure supplement 2**). Similarly, this represents a small reduction relative to the MAE (1.07 kcal) produced by predicting the dataset average affinity for each datapoint

The dopamine receptor also proved to be more challenging than σ_2_, particularly for AF3. With D4 with 205 confirmed actives (apparent *K_i_* values between 3.1 nM and 7.8 µM) and 336 inactive false positives with, we observed that actives were distinguished from docked false positives by docking (−63.9 vs −52.0 kcal/mol) and Boltz-2 (6.15 vs 5.72) scores, both with AUC ∼ 71. In contrast, AF3 L-pLDDT values showed little separation (65.4 vs 66.2) and an AUC = 46.4, which is slightly worse than random (**Fig. 5**c). There was also no clear evidence of early ligand enrichment with co-folding methods over docking methods from semilogarithmic ROC plots across the three virtual screen targets (**Figure 5—figure supplement 3**). Among active molecules, the docking correlation was slightly stronger for D4 than it was for σ_2_ (r = −0.247), but performance degraded for AF3 (r = −0.109, notably a positive correlation is expected) and Boltz-2 pIC_50_ (r = 0.284). Despite this reduced linear correlation for the Boltz-2 pIC_50_, the MAE was 0.89 kcal (0.55 kcal after correction). The performance degraded to 1.15 kcal (0.57 kcal after linear correction) under the optimistic assumption of 2X IC_50_ for all false positives and to 1.97 kcal (1.4 kcal after linear correction) under the randomly assigned assumption (**Figure 5—figure supplement 2**). This result is also a small improvement over the MAE (1.46 kcal) produced by predicting the dataset average affinity for each datapoint

An alternative way of looking at how well the scores can identify active ligands is to perform a hit enrichment analysis using normalized ranking scores (pProp, the negative logarithm of the percentage rank (Liu et al. 2025): a molecule that ranks among the top 0.1% of the docked library would have a pProp of 3, one that ranked among the top 0.01% of the docked library would have a pProp of 4, and so forth). pProp assigned here is based on the ranking within the docked list, for direct comparison between scoring methods and largely confirmed the trends above (**Figure 5—figure supplement 4**). Both DOCK3.7 and Boltz-2 clearly enriched actives across most targets, with hit rates plateauing at 40 - 50% for high-ranking compounds. AF3 L-pLDDT showed meaningful and comparable enrichment only for AmpC, while providing no enrichment signal for σ_2_ and D4. Overall, the success of the co-folding methods at predicting structures and correlating with affinities among true Mac1 inhibitors was far less apparent in their ability to distinguish true-from false-positives among docking hit-lists, and to correlate with affinities among these highly diverse molecules. Indeed, the methods showed little advantage over docking, a method that is about four orders of magnitude faster. An important caveat is that the hit-lists were composed of molecules prioritized by docking in the first place, giving it an advantage on these particular sets.

## Discussion

A striking observation from this study was the ability of the co-folding methods to predict ligand structures across 551 previously unpublished Mac1 complexes, and for the co-folding confidence metrics to correlate with ligand affinities. By both criteria all co-folding methods were meaningfully better than docking, at least as represented by DOCK3.7. Affinity prediction by Boltz-2 also performed well with mean absolute error that was ∼0.2kcal better than a baseline expectation of simply using the mean IC_50_ value. This small improvement was a trend that also held up in the large docking sets as well. Conversely, when confronted with high-anking molecules from large-library docking, a diverse mix of true- and false-positives with thousands of novel molecules prospectively tested against three unrelated targets, co-folding was not meaningfully better than docking at ligand categorization or at affinity prediction, indeed often worse. This despite the four log-orders greater speed of the docking and its well-known approximations and errors, long a topic of regret in the community (Jorgensen and Tirado-Rives 2005; Irwin and Shoichet 2016). These apparently contradictory results may be reconciled by the different tasks of predicting geometries for molecules that were all true ligands versus that of categorizing binders and non-binders within a highly diverse docking hit-list intentionally dissimilar to known ligands.

It may be that success and relative failure in these different tasks points also to a complementarity between physics-based docking and co-folding in the related, but different tasks of ligand discovery and optimization. The weak correlation between the methods (**Figure 5— figure supplement 5**) further suggests complementarity between deep learning and physics based approaches. Docking and related methods are designed for categorization among diverse, novel sets of candidate molecules, drawn from the vast new make-on-demand space (Gentile et al. 2020; Radaeva et al. 2023; Sadybekov et al. 2022; Sadybekov and Katritch 2023; Gorgulla et al. 2021; Lyu et al. 2019, 2023; Liu et al. 2024, 2025), whereas the co-folding methods, relying on learned patterns, may be relatively less effective. Where they have complementary strengths is in predicting the accurate geometries of ligands, once known, and their ranking within a series, both of which are crucial for affinity maturation once a series is discovered and with both of which docking has historically struggled. It is in these two linked but different aspects of ligand discovery and ligand optimization where the true complementarity between the methods may lie, at least for now.

Several caveats merit airing. The Mac1 benchmark, while diverse by ECPP4 Tc, is nevertheless dominated by four groups when clustered by MCS and benefits from a large training set of fragment molecules bound to the protein in the training set (Schuller et al. 2021). This may inflate apparent pose recovery rates, although Boltz-2 predicted poses were poorly recapitulated even with its much later training cutoff dates. Unexpectedly, even when the co-folding methods got important conformations of the enzyme binding site wrong, the ligands were still often posed correctly, speaking to a non-physical aspect of the ML methods that others have also described (Masters et al. 2025; Škrinjar et al. 2025). Even though the prospective docking hit lists included thousands of diverse molecules, they still represent only three targets, and so offer a limited view of molecular recognition. Finally, comparing co-folding to docking based on hit-lists themselves selected by docking is arguably unfair to co-folding. Counterbalancing this is the inclusion, in each of the three hit lists, of molecules that had mediocre and poor docking scores intentionally selected to test the correlation between docking score and hit-rate. Here too, the correlation between co-folding score and likelihood to bind, what we sometimes call a “dock-response-curve” was no better than docking’s, often worse (**Figure 5—figure supplement 4**).

These caveats should not obscure the central lessons from this study. Against a benchmark of 551 topologically diverse Mac1 ligand-bound structures, determined to true atomic resolution and largely dissimilar to structures already in the PDB, each of the co-folding methods predicted ligand poses and interactions with high-fidelity to the experimental structure. For Boltz-2, there was a strong and highly significant correlation between predicted affinity and experimental IC_50_ (compared to a modest correlation for AF3 L-pLDDT and experimental IC_50_). In both cases, co-folding performance was much better than that of DOCK3.7, although the correlation was only significantly different for Boltz-2 pIC_50_ versus experimental pIC_50_. Conversely, when the more homogeneous benchmark of true ligands was replaced by a diverse set of docked molecules, composed of what would turn out to be hundreds of true ligands and thousands of false positives, the co-folding methods did no better than docking and often did worse. These observations are supported by a fascinating study on some of the same ligand sets as investigated here, using AlphaFold3, reaching similar conclusions (Menon et al. 2025). The relatively strong performance by docking is surprising given its well-known weaknesses and its much reduced computational cost (and therefore greater speed). What emerges are approaches that may be complementary in their roles in ligand discovery (docking) and structure-based ligand optimization (co-folding), areas that will merit future research and application.

## Methods

### Mac1 X-ray crystallography

Ligands were soaked into SARS-CoV-2 Mac1 crystals (P4_3_ crystal form (Schuller et al. 2021)), and data collected and reduced, as described previously (Gahbauer et al. 2023; Correy et al. 2025). We modeled ligands into PanDDA event maps (Pearce et al. 2017) using ligand coordinates and restraints generated with phenix.elbow (Moriarty et al. 2009).

### Ligand Similarity, Chemical Diversity and Memorization Analyses

To test whether pose-prediction success reflected latent memorization, we quantified chemical similarity between test and training ligands using two complementary metrics. First, extended-connectivity fingerprints (ECFP4) were computed for all molecules, and Tanimoto coefficients (TC) were used to identify, for each test ligand, the most similar compound present in the training corpus (Chai-1 entries deposited before 1 December 2021 and AlphaFold3 entries before 30 September 2021). The training date cutoff for Boltz-2 is 1 June 2023 (**Supplementary File 6).**

As another measure of structural similarity, the maximum common substructure percentage (MCS%) was calculated using the rdFMCS algorithm in RDKit, reporting the proportion of heavy atoms shared between two ligands. To evaluate chemical diversity within benchmark datasets, pairwise TC or MCS% values were computed across all ligands. Best-First Clustering (BFC) based on these similarity metrics was used to estimate the number of unique chemotypes represented in each dataset. Principal component analysis (PCA) was used to visualize the resulting chemical space, and to identify an appropriate threshold for MCS-based scaffold grouping. Within the Mac1 benchmark set, applying an MCS cutoff of 35% yielded four distinct scaffolds, in agreement with PCA-derived cluster separation. The same clustering threshold was subsequently applied to define scaffold families for the AmpC, σ_2_ and D4 benchmark datasets.

### Molecular Docking of Mac1 compounds

The modelled orthosteric site of the Mac1 model was available from the PanDDA analysis group deposition in the Protein Data Bank (PDB), with PDB ID: 5SQW (Gahbauer et al. 2023). We used this structure because the inhibitor (Z5014193706) was the most potent molecule with a structure determined around the same time as the ligands in this dataset were tested. This structure was used to generate matching spheres, which are later used by DOCK3.7 to pre-sample ligand conformations into binding sites by superimposing ligand atoms on spheres in the site. 45 matching spheres were used, and the docked ligands were scored by summing receptor-ligand electrostatic and van der Waals energies and corrected for context-dependent ligand desolvation. Receptor structures were protonated using Reduce. Partial charges from the united-atom AMBER force field were used for all receptor atoms. Potential energy grids for the different energy terms of the scoring function were precalculated using AMBER for the van der Waals term and the Poisson-Boltzmann method QNIFFT for electrostatics. Context-dependent ligand desolvation was calculated from the Generalized Born method. Ligands were protonated at pH7.4 (ChemAxon), and each protomer was rendered into 3D using Corina, followed by conformational sampling using Omega. For the docking campaign of 551 novel compounds, each molecule was sampled in about 2,719 orientations and, on average, 105 conformations by. Overall, over 290 million complexes were sampled and scored.

### Co-folding

All three co-folding methods (AlphaFold3, Chai-1 and Boltz-2) were available from following sources:

- AlphaFold3: https://github.com/google-deepmind/alphafold3
- Chai-1: https://github.com/chaidiscovery/chai-lab.git
- Boltz-2: https://github.com/jwohlwend/boltz

Ligand SMILES and amino acid sequences for the target protein were used to perform co-folding in an array of NVIDIA A40 GPUs. Multiple Sequence Alignment (MSA) was performed with the Jackhammer module. Co-folding tools will generate predicted protein-ligand complex poses (mmCIF files), and different confidence metrics for these predictions. Chai-1 and AlphaFold3 generated global confidence scores (pTM, ipTM), whilst AlphaFold3 had local atom-specific metrics, such as, pLDDT, PAE and contact probabilities in large vectors. To put an emphasis on prediction of ligands, we computed a ligand-centric pLDDT (L-pLDDT) by taking an arithmetic mean of pLDDT on ligand atoms, and a ligand-centric PAE (L-PAE) by taking an average of all PAE elements that contain ligand atoms. Minimum PAE (mPAE) was obtained by scanning all PAE elements that involve ligand atoms, and finding the smallest error. Boltz-2 generated all the confidence metrics in Chai-1 and AlphaFold3, but its affinity module generated an output of the binding likelihood (probability metric) and the predicted binding affinity (Boltz-2 pIC_50_) of compounds. Scripts to prepare input .json, .fasta files, postprocess output files, are available in https://github.com/jongbin99/Cofolding.

### Benchmarking Co-folding on Experimentally Validated Compounds

With novel compounds discovered by fragment-based screening, 551 Mac1 complexes were resolved by the crystallography work. The same sequences were used to perform co-folding with AlphaFold3, Chai-1 and Boltz-2. For further sets of analyses with docked hit lists on targets from the Large-Scale Docking (source: lsd.docking.org)(Hall et al. 2025), only AlphaFold3 and Boltz-2 were used. For each system, we redefined the threshold for true hits and non-binders as described herein:

- AmpC β-lactamase: Among 1.7 billion docked molecules, the top scoring 1% were selected (n = 1,293) (Liu et al. 2025). We selected ones with apparent Ki < 400 μM (n = 247) as true hits, and the rest were regarded as non-binders.
- σ_2_: True hits among these docked molecules were previously defined as displacing greater than 50% of 3H ditolylguanidine ([3H]DTG) to σ_2_ (Alon et al. 2021), but in our work, the cutoff is adjusted to 25% displacement, giving a less stringent threshold to include more molecules into the space of true hits. Single concentration point measurement (at 1 μM) is converted to an apparent Ki value using Cheng-Prusoff equation 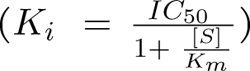, with pre-calculated values of 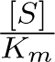based on radioligand displacement (Alon et al. 2021). This yields a non-binding cutoff of an apparent Ki < 2.0 μM.
- Dopamine D4: For the dopamine receptor D4, 138 million molecules were docked and then prioritized synthesizable high-ranking docked molecules (n = 549). True hits from these molecules were defined as displacing more than 50% of 3H-N-methylspiperone (Lyu et al. 2019), but similarly to σ_2_, we adjusted the cutoff to 25% displacement. Apparent Ki values are calculated similarly to σ_2_ case, but at 10 μM. This yields a non-binding cutoff of an apparent Ki < 7.8 μM.

### Ligand Pose Evaluation via RMSD against crystal structures

Predicted ligand binding modes were benchmarked against experimentally determined reference poses from ultra-high-resolution (∼1.0 Å) crystal structures. The Mac1 protein receptor was first superimposed by least-squares fitting of all Cα atoms using PyMol. The heavy-atom root-mean-square deviation (RMSD) between the aligned co-folded ligand and its crystallographic counterpart was then calculated. Before RMSD evaluation, both predicted and reference ligands were confirmed using a uniform RDKit-based sanitization pipeline (Buttenschoen et al. 2024). SMILES strings were converted to RDKit Mol objects, chemically validated (valence, aromaticity), and assigned bond orders from a canonical SMILES-derived template. We then check that ground-truth (experimental) and co-folded ligand coordinates for all molecules to have valid chemistry as described with the SMILES-derived template. A substructure match then enabled a one-to-one correspondence of heavy atoms between the ground-truth and co-folded poses for comparison. The crystal ligand and co-folded ligand had the center-of-mass (COM) calculated separately, and measured the distance between each COM. If the distance exceeds 2.5 Å, we would expect the RMSD to be large, indicating that ligand poses are not modelled accurately in an orthosteric pocket (Nittinger et al. 2025). For all the predicted ligand poses, rdMolAlign.CalcRMS from Rdkit.Chem is used for calculating the ligand RMSD between co-folding and the ground-truth crystal pose.

### Hit-rate curves for different scoring methods

To get hit-rate curves for co-folding and docking scores across three targets of docked hitlists (AmpC, σ2, D4), molecules in the list are ordered by increasing score (for DOCK) and decreasing score (for L-pLDDT and Boltz-2 pIC_50_). A rolling window of 100 was used for AmpC and σ_2_, and 50 for D4 (Liu et al. 2025), calculating the hit rate as a fraction of molecules with experimental binding affinity that match our definition for true hits described above (refer to Benchmarking Co-Folding on Experimentally Validated Compounds in Method section). The docked hit-list is a mixture of known actives and non-binders that include false positives based on docking scores. For direct comparison of hit classification performance between DOCK energies, AF3 L-pLDDT and Boltz-2 pIC_50_, the pProp (the negative base 10 logarithm of the fractional rank) was assigned within the list, instead of the billion-scale full docking list. For each window, we used a 95% Wilson confidence interval with normal approximation for the hit rate.

## Statistical Analysis

Pearson’s correlation coefficients were used throughout the paper to quantify linear associations between two variables in the study. We have also reported a mean absolute error of data points with respect to the regression line for Boltz-2 affinity pIC_50_ against the measured pIC_50_. The mean absolute errors are also calculated for the perfect fit (Pearson r = 1) and for the fixed measured pIC_50_ (Pearson r = 0), to give an indication of how accurate the Boltz-2 scores are with respect to actual binding energies (in kcal), in terms of correlations and absolute errors. Friedman and post-hoc Conover tests were used to compare how each of AF3, Boltz-2 and DOCK3.7 score predicts for actual binding affinity, and Holm-Bonferroni correction for pairwise comparison p-values (Ash et al. 2025). The distribution of scores for known actives and docked false positives were compared using the Kolmogorov-Smirnov (KS) non-parametric statistical test, as shown in violin plots, and the KS statistic value indicates how distant the two distributions are, and p-values show the extent of statistically detectable differences. All analyses were performed using Python packages.

## Supporting information

Supplementary File 1 - X-ray data collection statistics and PDB accession codes

Supplementary File 6 - PDB accessions by training-set cutoff date

Supplementary File 2 - Pairwise ligand similarity by ECFP-4 Tc and MCS%

Supplementary File 3 - Mac1 ligands with co-folded and docked poses and scores

Supplementary File 4 - Ligand interactions in ground-truth and AF3 models

Supplementary File 5 - Docked hits for AmpC, sigma2 and D4

## Acknowledgements

We acknowledge funding from NIH GM122481 (to B.K.S.), GM145238 (to J.S.F.), and U19AI171110 (to N.J.K). We thank Pat Walters for helpful comments. The synchrotron X-ray diffraction data used to determine Mac1 structures were collected at beamline 8.3.1 of the Advanced Light Source (ALS) and beamlines 9-2, 12-1 and 12-2 of the Stanford Synchrotron Radiation Lightsource (SSRL). The ALS, a U.S. DOE Office of Science User Facility under contract no. DE-AC02-05CH11231, is supported in part by the ALS-ENABLE program funded by the NIH, National Institute of General Medical Sciences, grant P30 GM124169. Use of the SSRL, SLAC National Accelerator Laboratory, is supported by the U.S. Department of Energy, Office of Science, Office of Basic Energy Sciences under Contract No. DE-AC02-76SF00515. The SSRL Structural Molecular Biology Program is supported by the DOE Office of Biological and Environmental Research, and by the National Institutes of Health, National Institute of General Medical Sciences (P30GM133894).

## Author contributions

Cofolding: J.K., B.W.H., R.R.

X-ray crystallography: G.J.C.

HTRF assay: Y.U.D., M.G.V.S., M.E.D.

Compound design: G.J.C., M.M.R., T.T., R.L.G., P.J., R.J.N., E.R.H., A.R.R.

Compound synthesis: T.T., R.L.G., P.J.

Docking: M.M.R.

Data analysis: J.K., B.W.H., O.M.

Supervision and project management: S.G.

Supervision and manuscript editing: N.J.K., A.A.

Writing and editing: J.K., G.J.C., M.M.R., A.R.R., A.A., B.K.S., J.S.F.

## Data availability

Coordinates and structure factors for the Mac1 ligand complexes determined in this work have been deposited in the Protein Data Bank. The 468 accession codes are listed in Supplementary File 1, together with the data collection and refinement statistics for each structure. A further 22 Mac1 complexes analysed here were deposited as part of earlier studies. Their accession codes and source publications are listed in the same file. The Mac1 structure used to build the docking receptor is PDB 5SQW. Co-folded and docked poses, confidence metrics and scores for every ligand are provided in Supplementary File 3. The experimentally tested compounds and measured potencies for the AmpC β-lactamase, σ2 and dopamine D4 docking campaigns are provided in Supplementary File 5. These data were taken from previously published screens (Lyu et al. 2019; Alon et al. 2021; Liu et al. 2025), and the corresponding hit lists are available from lsd.docking.org. Scripts used to prepare the co-folding inputs and to analyse the resulting poses and scores are available at https://github.com/jongbin99/Cofolding.

## Supplementary Information

**Figure 1—figure supplement 1.**
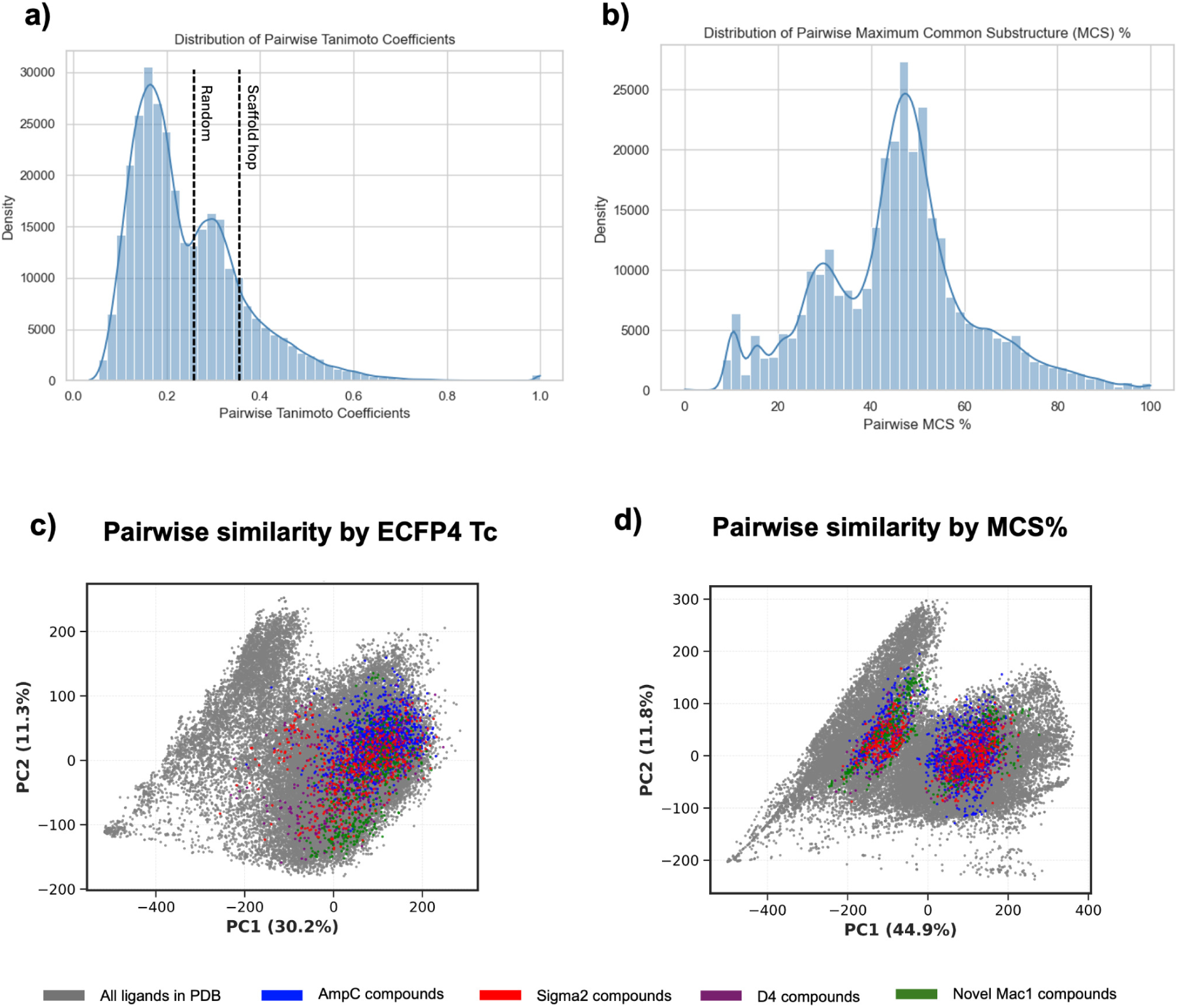
Distribution of pairwise similarity for Mac1 ligands and an overlay of clusters of compounds tested on entire ligand set deposited in PDB. Pairwise similarity of Mac1 compounds by a) ECFP-4 Tanimoto Coefficients (TC) with lines indicating values for a ‘scaffold-hop’ (Tc of 0.35), or ‘random’ (Tc of 0.25), b) Maximum Common Substructures (MCS%). Principal Component Analysis of all the ligands deposited in PDB, and the compounds tested for co-folding, to show the diversity of the compounds chosen for the study over the entire ligand space of the training set, by c) ECFP-4 Tc and d) MCS%.

**Figure 1—figure supplement 2.**
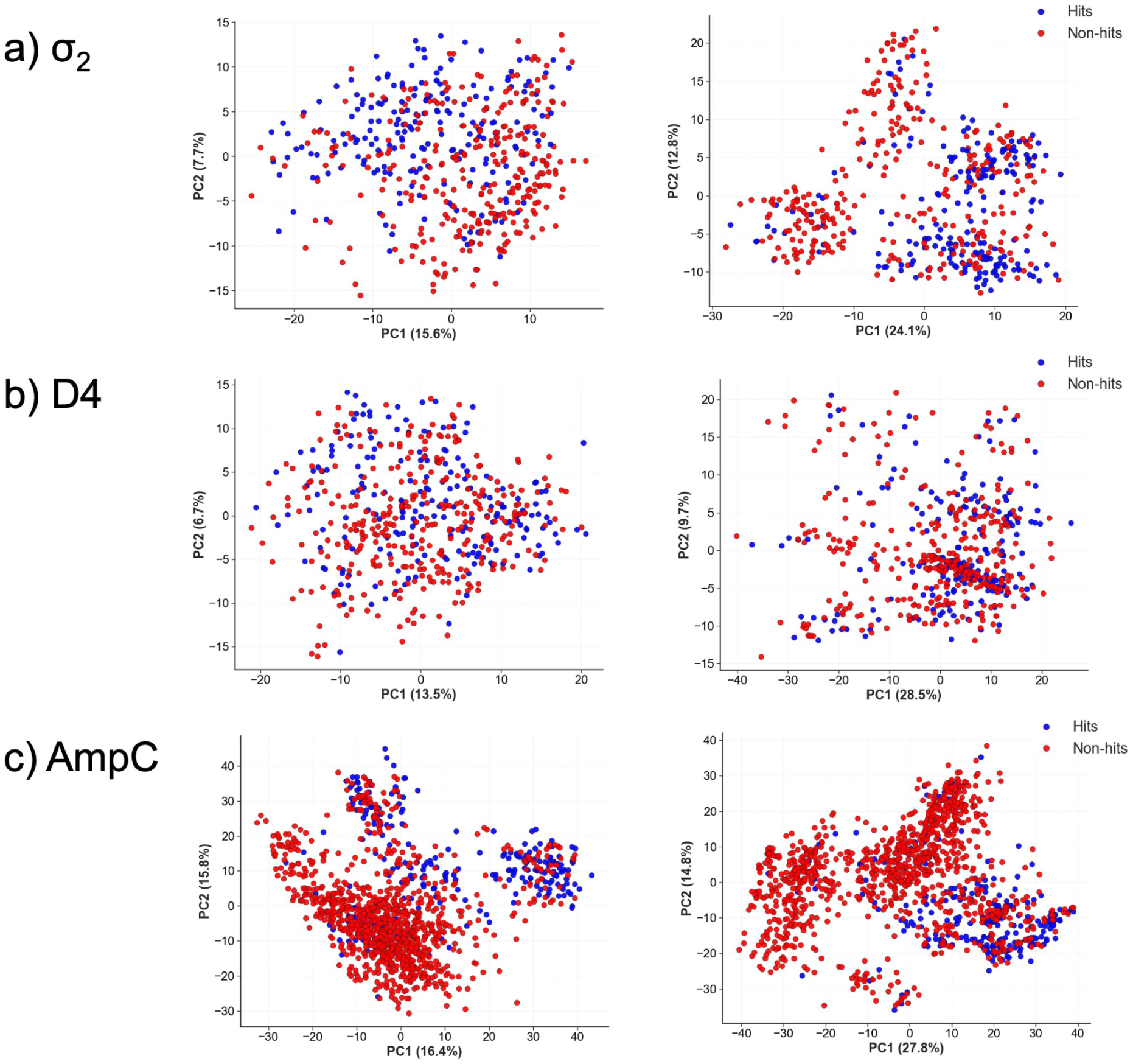
Benchmarked datasets are clustered based on TC and MCS% to predict the number of scaffold series. Principal Component Analysis (PCA) plots make pairwise comparisons of molecules in the docked hit lists of a) σ_2_, b) D4, c) AmpC for similarity using two different metrics: TC (first column) and MCS% (second column). Red points indicate docked false positives in the hit list, and blue points indicate known actives. Mac1 clustering plots were shown in Fig.1 and an asterisk indicates we only have known ligands for the Mac1 set. The number of scaffolds based on cluster heads are determined by either TC > 0.35, or MCS > 35%. The table gives the summary of the clustering analysis of the datasets used.

**Figure 2—figure supplement 1.**
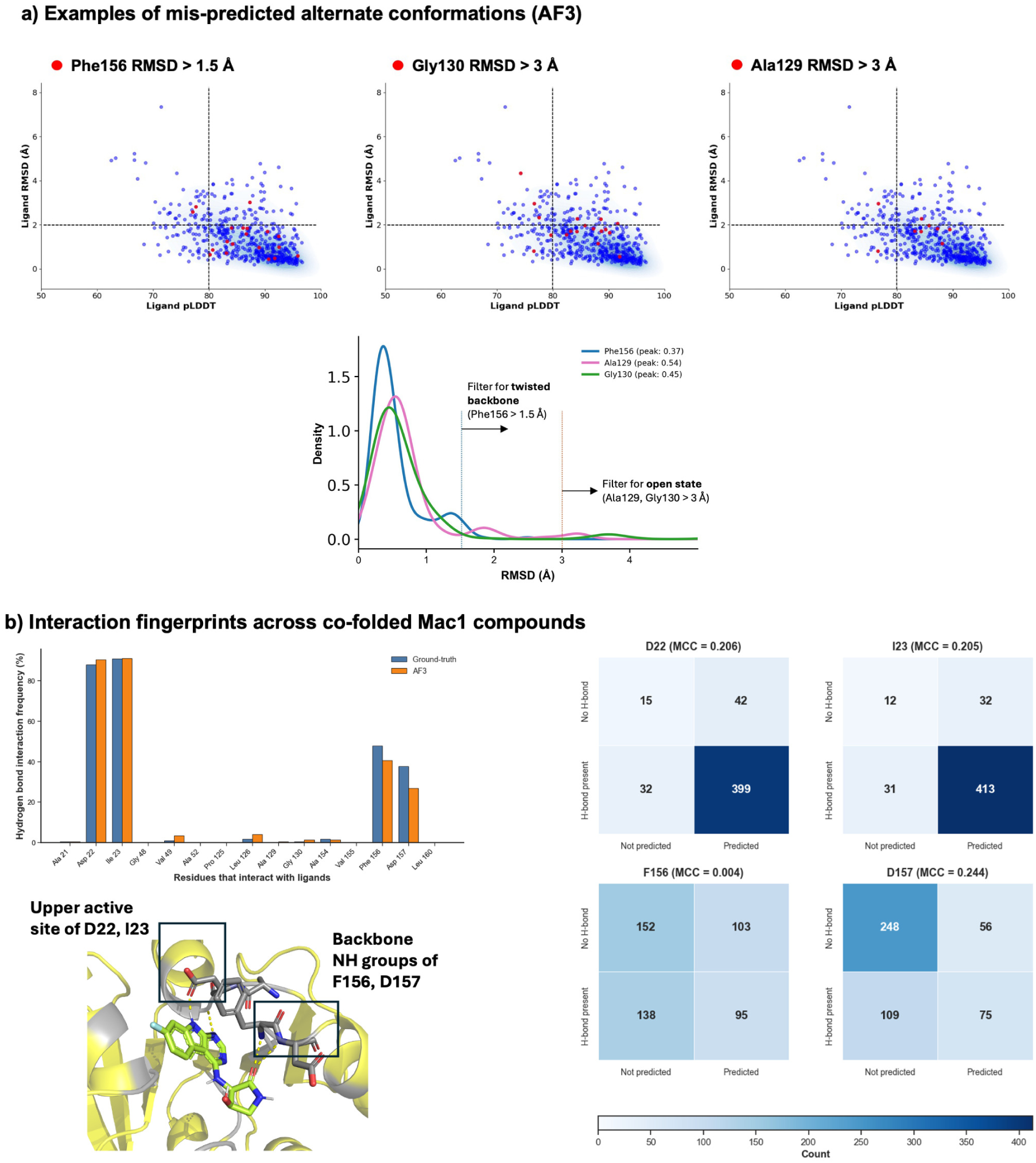
Detailed predictions of alternate residue conformations and interactions in co-folded poses. **a)** Mis-prediction of alternate conformations of the binding pocket, including twisted backbone (Phe156 RMSD > 1.5 Å) and open structure (Gly130 RMSD, Ala129 RMSD > 3 Å) did not produce false positive or false negative pose prediction. The density distribution plot shows RMSD between residues in PDB ID: 5SQW, and 489 complexes obtained from crystallography, a filter that was used to define alternate conformations. b) Hydrogen bonds between ligands and residues within 5 Å were counted for crystal, and for AF3 co-folded structures. Interactions shown with 4 hotspot residues (D22, I23, F156, D157) were compared and confusion matrices show absolute counts of True positives, True negatives, False positives and False negatives. Matthews Correlation Coefficients are calculated for each residue.

**Figure 3—figure supplement 1.**
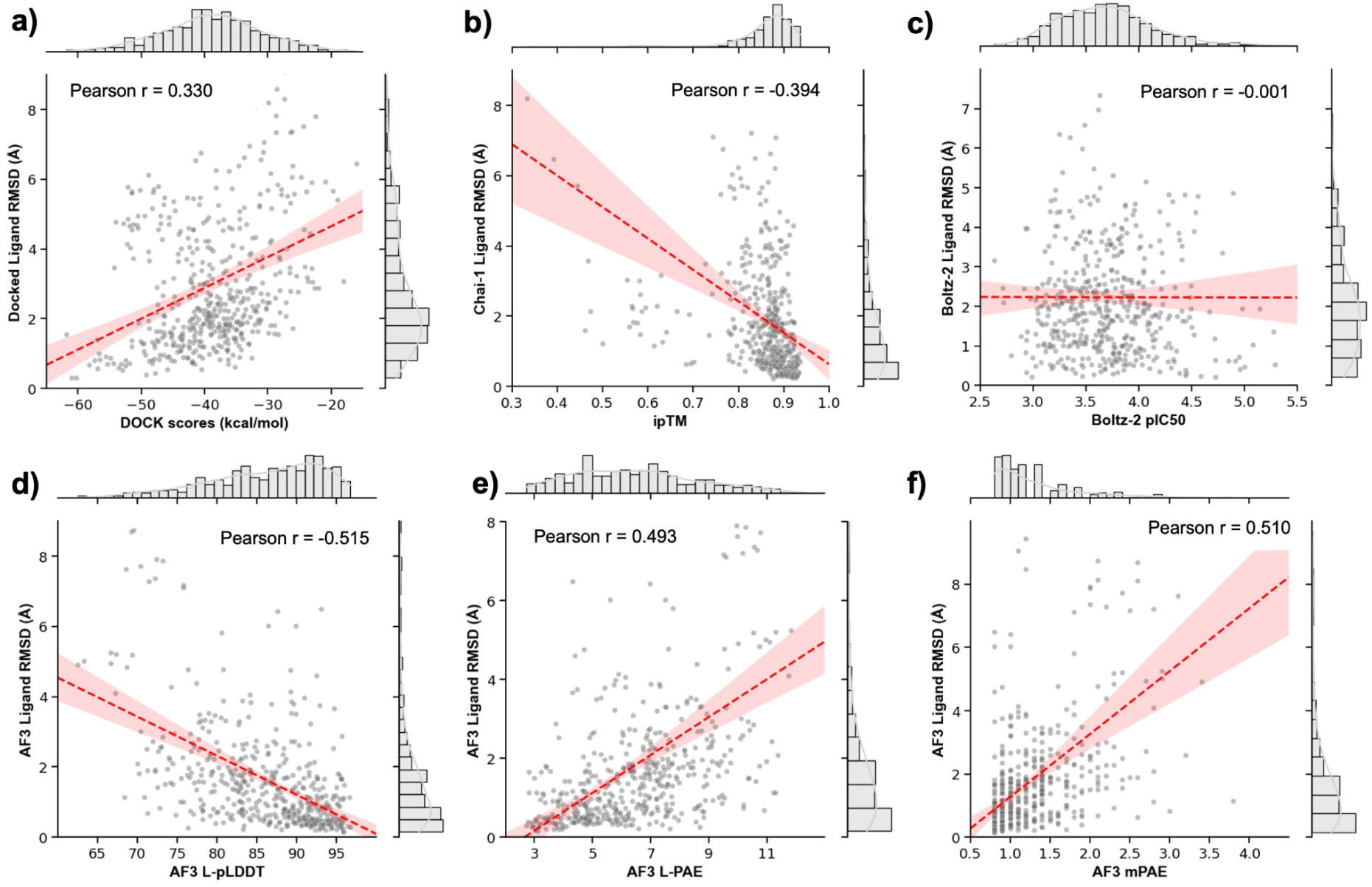
Correlations between Mac1 pose recovery with co-folding/docking scoring metrics. **a)** Docked pose RMSD is compared with DOCK scores (in kcal/mol), b) Chai-1 RMSD is compared with interface predictive TM-Score (ipTM), and c) Boltz-2 pose RMSD is compared with predicted Boltz-2 pIC_50_ affinity score. AF3 RMSD is compared with three possible scoring metrics from AF3: d) Ligand-specific pLDDT (L-pLDDT), e) Ligand-specific PAE (L-PAE), f) minimum PAE (mPAE).

**Figure 3—figure supplement 2.**
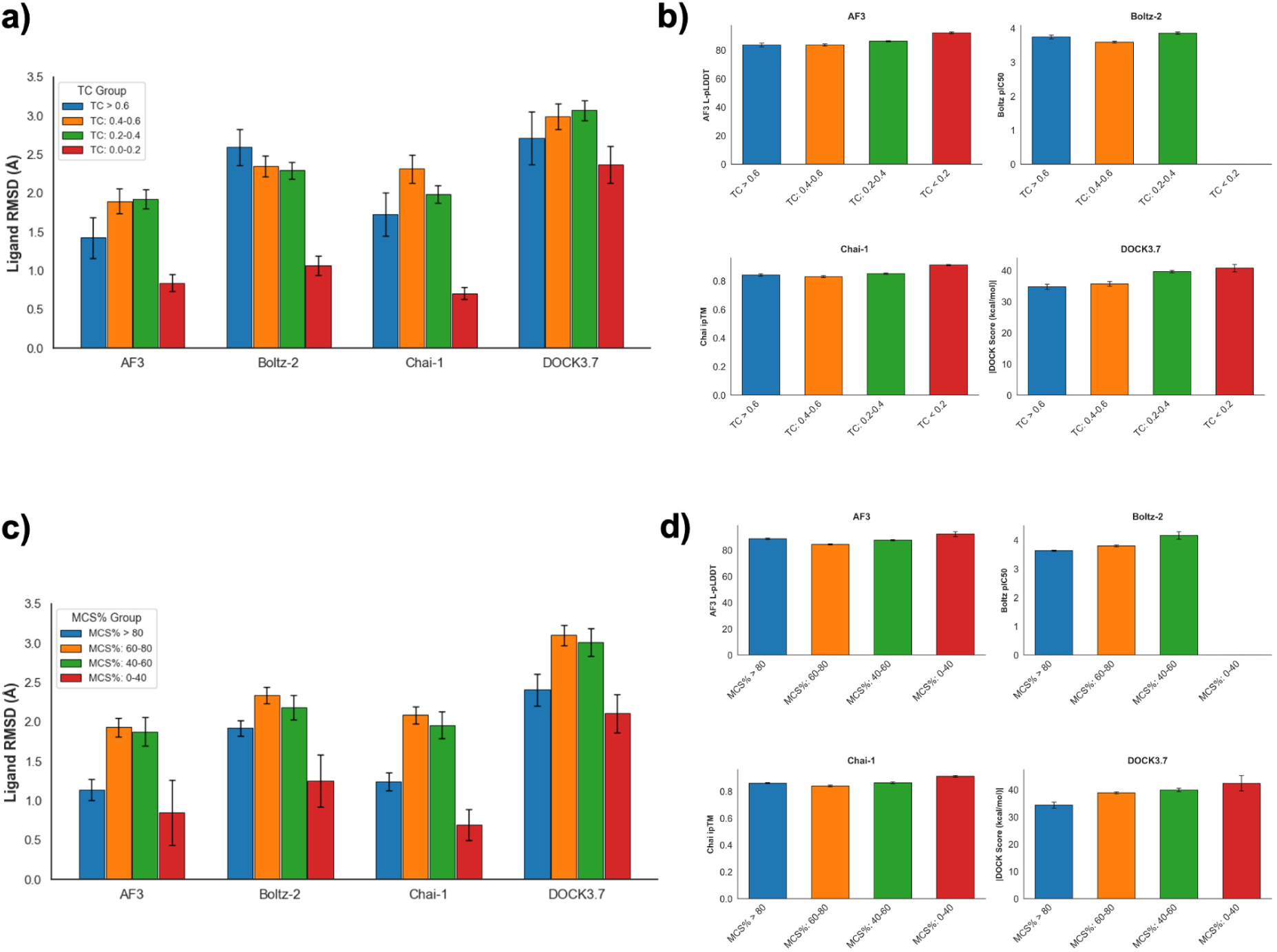
Pose accuracy and co-folding, docking scores are compared for different similarity bins. **a)** Ligand RMSD between co-folded poses and docked poses are compared to the ground-truth for different Tanimoto Coefficient (TC) bins (< 0.2, 0.2-0.4, 0.4-0.6, > 0.6), b) Scores for each method (AF3 L-pLDDT, Chai-1 ipTM, Boltz-2 pIC_50_, DOCK3.7 energies) are compared for different TC bins. c) Ligand RMSD for different Maximum Common substructure (MCS%) bins (< 40, 40-60, 60-80, > 80), d) Scores for each method are compared for different MCS% bins. TC and MCS% of new molecules to the training set were calculated for each model, and for DOCK3.7 (the only non-co-folding method), the similarity was calculated against the AF3 training set.

**Figure 3—figure supplement 3.**
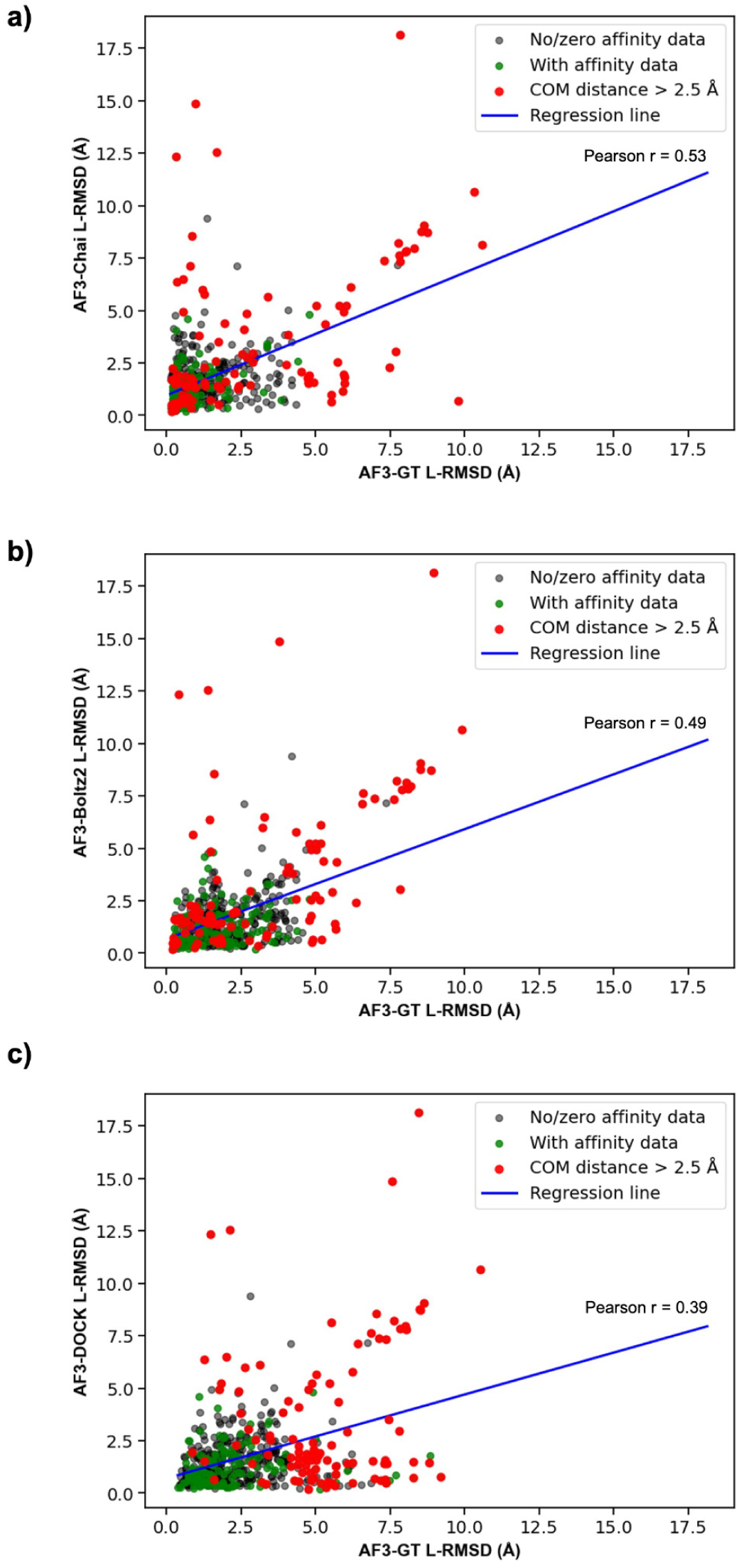
Correlation between AF3 pose-prediction error and errors from other co-folding and docking methods for 551 Mac1 compounds. **a)** AF3-Ground truth L-RMSD vs AF3-Chai L-RMSD, b) AF3-Ground truth L-RMSD vs AF3-Boltz 2 L-RMSD, c) AF3-Ground truth L-RMSD vs AF3-DOCK L-RMSD. Ligands with affinity data (n= 202) are marked in green, those with high COM distance are marked in red, and molecules with no affinity data are marked as grey. Pearson correlation coefficients and regression lines are shown.

**Figure 3—figure supplement 4.**
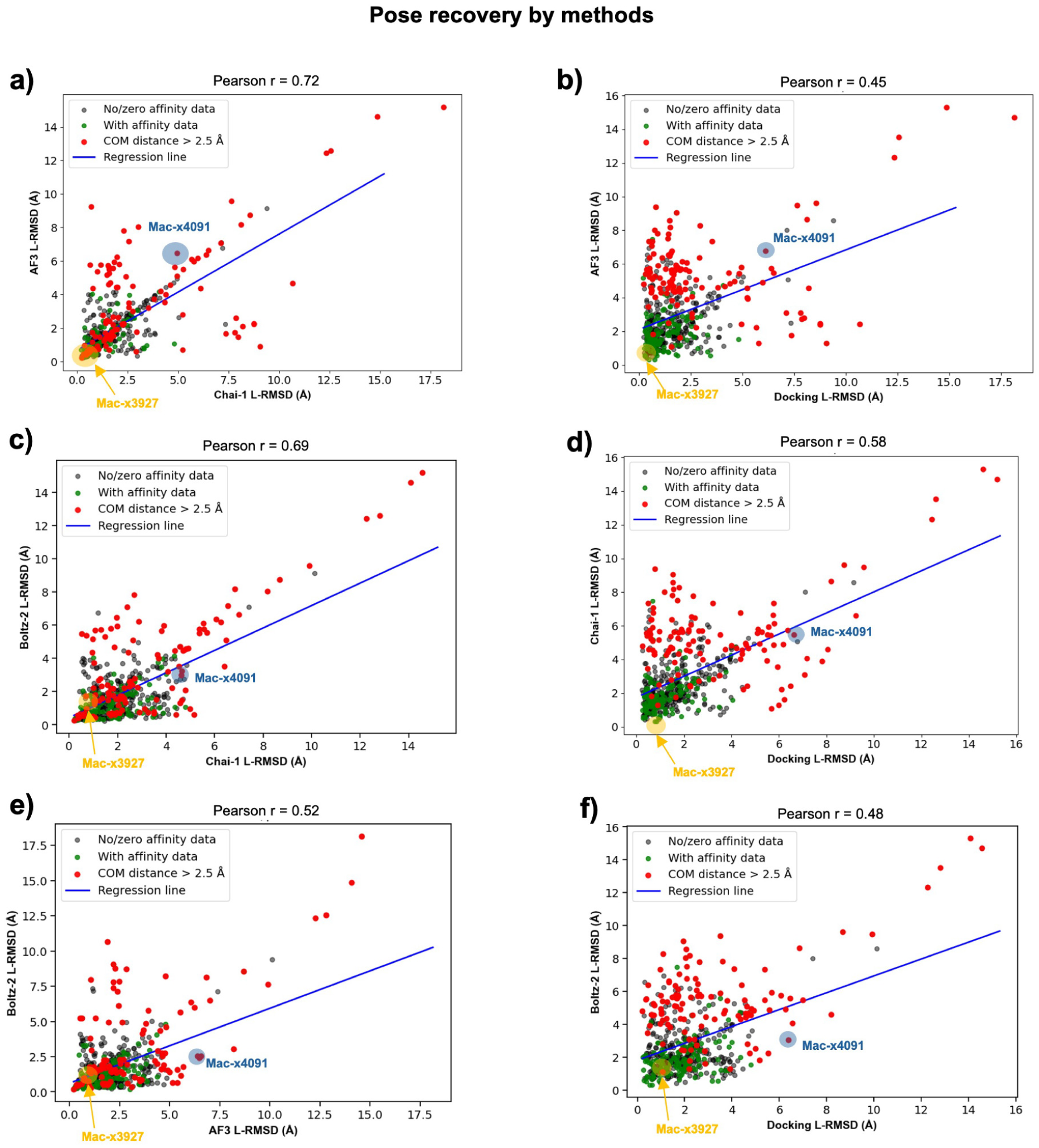
Pose recovery by methods of co-folding and docking against the ground-truth pose. **a)** AF3 vs. Chai-1, b) AF3 vs. DOCK3.7, c) Boltz-2 vs. Chai-1, d) Chai-1 vs. DOCK3.7, e) Boltz-2 vs. AF3, f) Boltz-2 vs. DOCK3.7. Molecules indicated (Mac-x4091, Mac-x3927) are exemplary ligands highlighted in Fig. 3, and L-RMSD values quoted here are all compared against the ground-truth pose.

**Figure 4—figure supplement 1.**
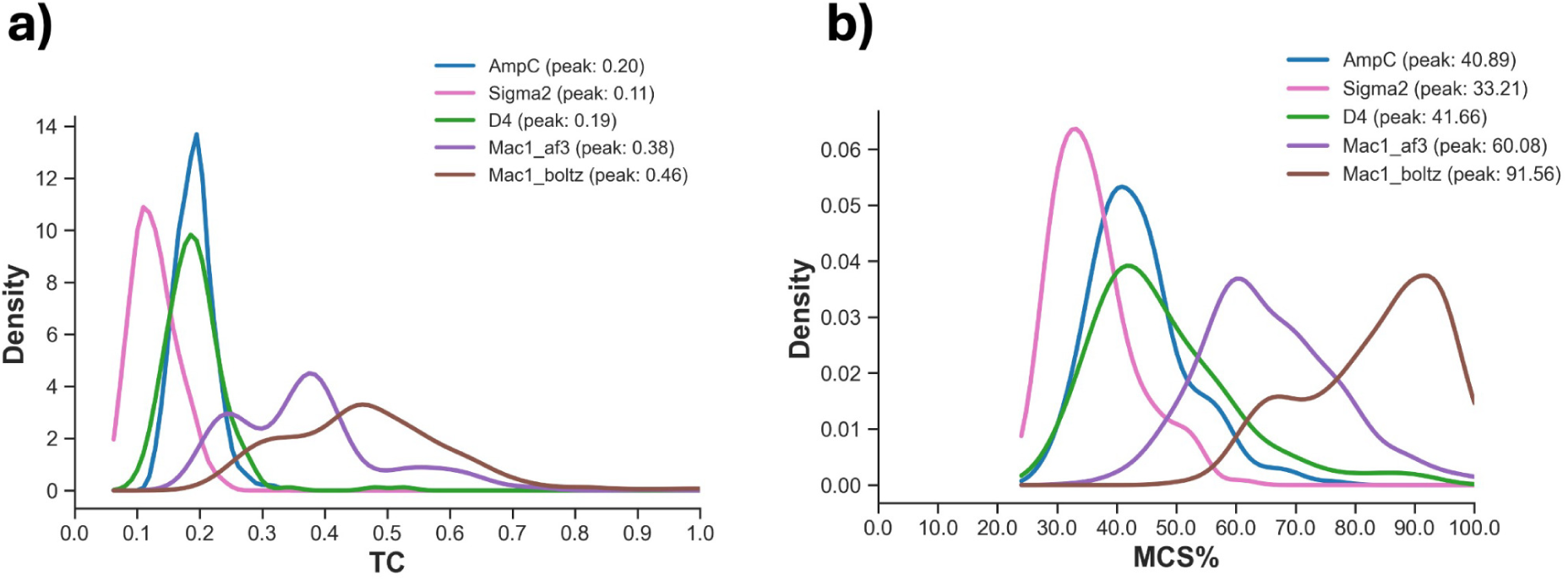
Actives from the benchmarked datasets are compared with resolved structures used to train models like AF3 and Chai-1. Actives from AmpC (n = 247), D4 (n = 205), σ_2_ (n = 201), Mac1 (n = 551) were compared with corresponding training sets, to give an indication how similar the actives were to those trained by co-folding models, based on a) Tanimoto coefficients, and b) Maximum Common Substructures (%). AF3 and Chai-1 have similar cutoff dates (indicated as Mac1_af3), but Boltz-2 has been trained with more recent PDB and is labelled as Mac1_boltz. Exact PDB IDs used for each model system is shown in Supplementary File 6.

**Figure 5—figure supplement 1.**
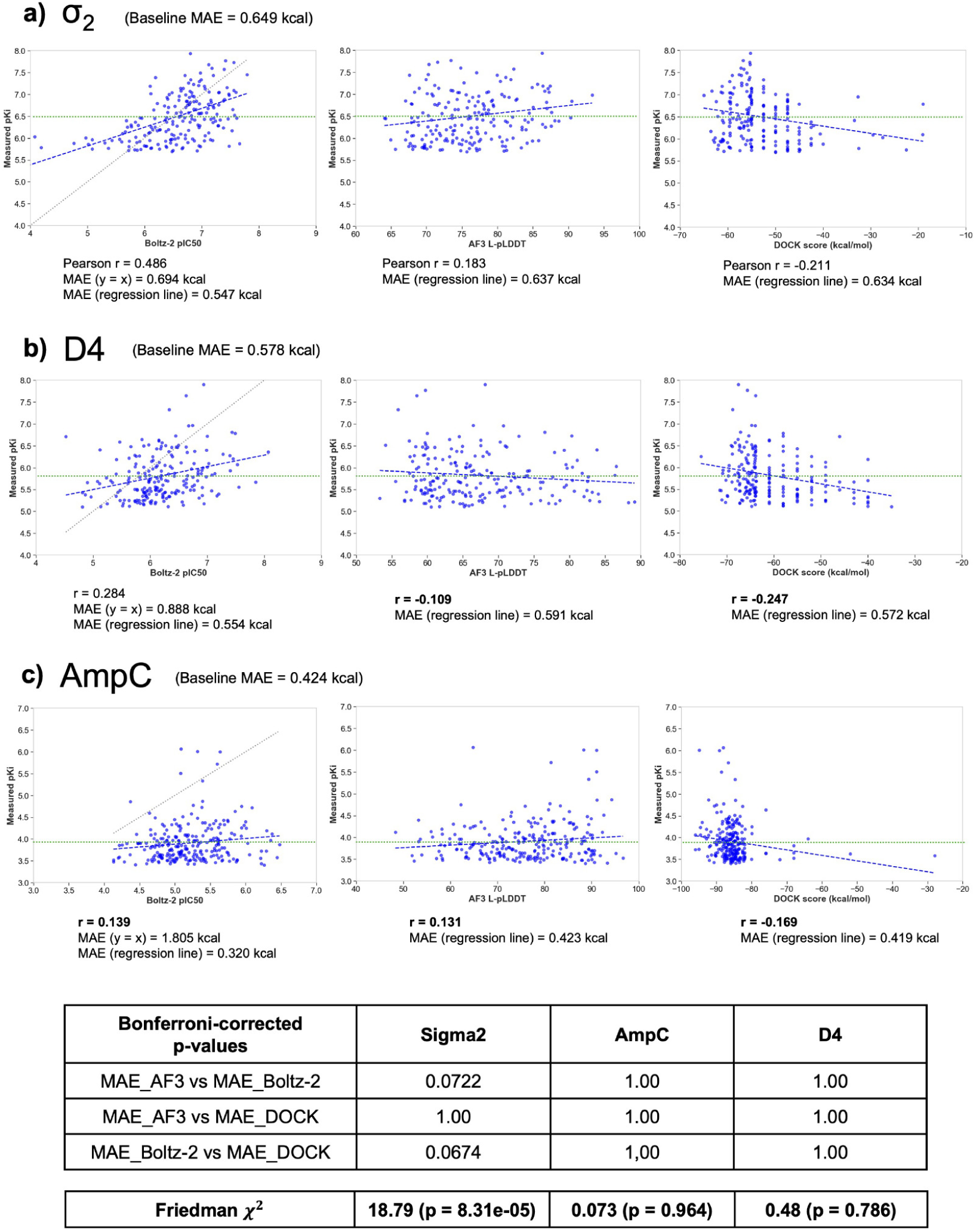
Discriminative power of co-folding and docking scores by correlation against experimentally measured *pK_i_* across different targets: a) σ_2_ (n = 201 actives), b) D4 (n = 205 actives), c) AmpC (n = 247 actives). The leftmost panel is for Boltz-2 pIC_50_ affinities, the middle panel for AF3 L-pLDDT and the rightmost panel for DOCK scores. All these scores are compared against measured *pK_i_* values. Mean Absolute Errors and Pearson correlation coefficients are calculated. Blue lines show the corrected regression line, black dotted line before the correction, and green horizontal line shows the baseline at an average measured pKi value. Friedman test with Conover post-hoc comparisons and Bonferroni corrections compare MAE between methods for each target system.

**Figure 5—figure supplement 2.**
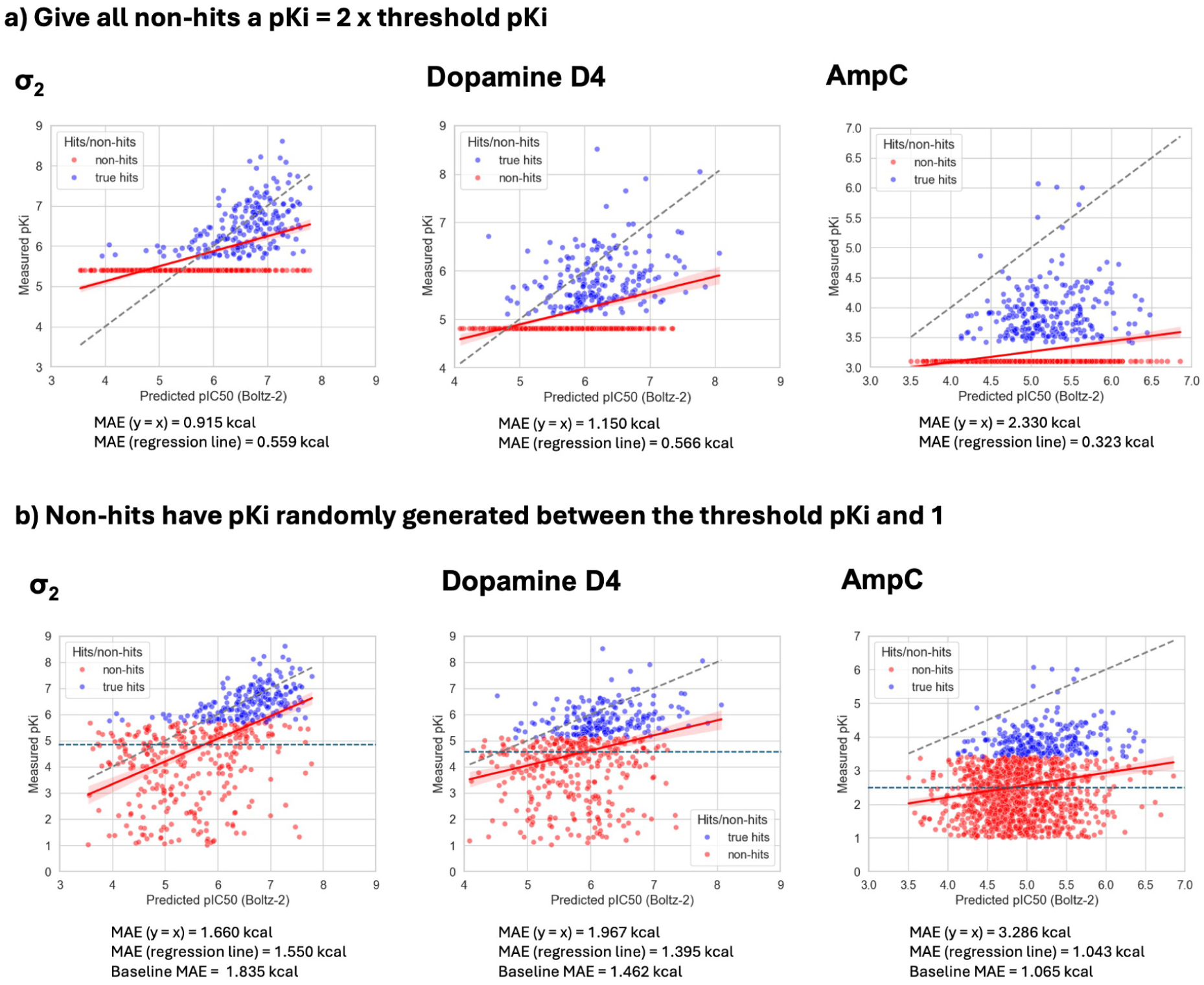
Different methods of treating randomly assigned *pK_i_* values for docked false positives in σ_2_, D4, AmpC docked lists. Mean absolute errors (before linear correction is shown in grey dotted line, after linear correction is the red regression line and blue line indicates the baseline when we predict all values at measured *pK_i_*) between the measured *pK_i_* and Boltz-2 pIC_50_ affinity scores when a) all non-hits are assigned a *pK_i_* = 2 x *pK_i,threshold_*; b) non-hits are randomly assigned a *pK_i_* value between the *pK_i,threshold_* and 1. Baseline MAE is only quoted for b), since the error could be misinterpreted when all non-binders’ *pK_i_* values are fixed.

**Figure 5—figure supplement 3.**
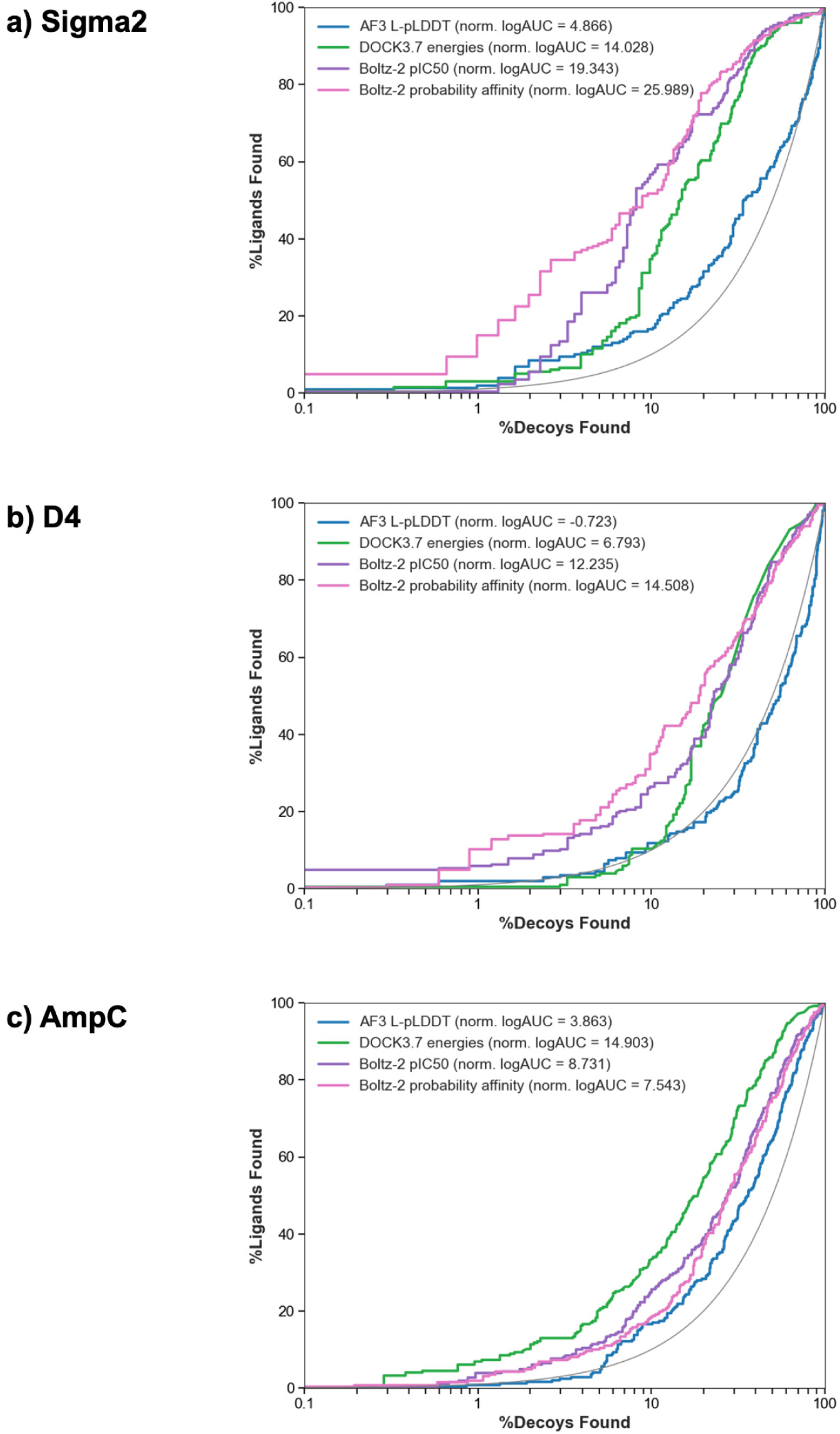
Semi-logarithmic ROC curves to differentiate actives over decoys using co-folding and docking scores for the three experimental benchmark datasets. Semi-logarithmic axes will consider early enrichment of ligands, using AF3 L-pLDDT, DOCK3 energies (kcal/mol), Boltz-2 pIC50 and Boltz-2 binary probability affinity values for a) Sigma2, b) Dopamine D4, c) AmpC β-lactamase.

**Figure 5—figure supplement 4.**
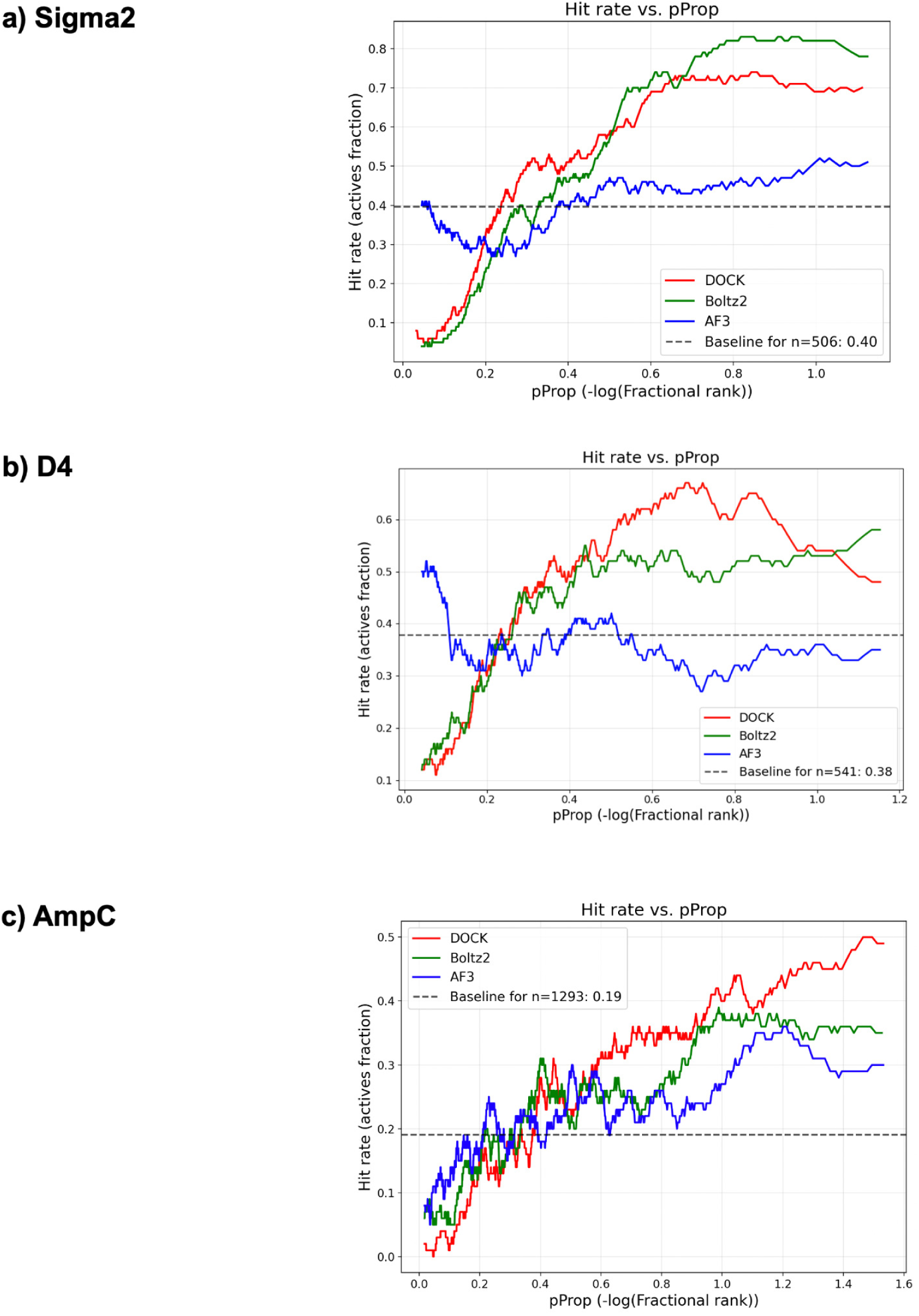
Hit rate curves for co-folding and docking scores for the three experimental benchmark datasets. Hit rate curve are plotted over a rolling window (window = 100 for AmpC, σ_2_ and 50 for D4) after ranking with pProp from the docked hit lists, from three different targets: a) σ_2_ (201 actives, 305 nonbinders), b) Dopamine D4 (205 actives, 336 non-binders), c) AmpC β-lactamase (247 actives, 1,046 non-binders).

**Figure 5—figure supplement 5.**
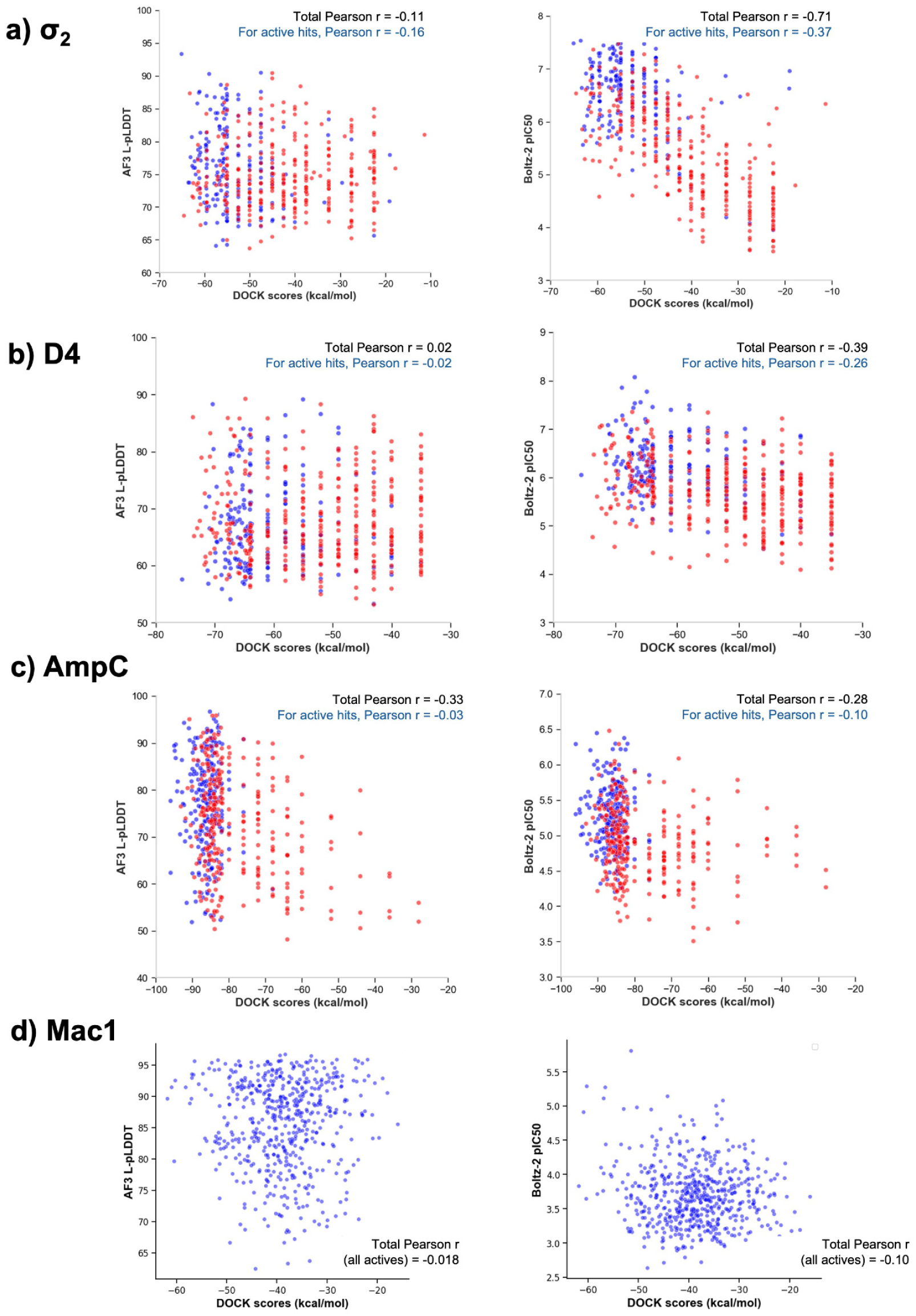
Correlation between DOCK scores and some co-folding scores (AF3 L-pLDDT, Boltz-2 pIC_50_). Blue points indicate known actives and red points are non-binders for each target system: a) σ_2_, b) D4, c) AmpC and d) Mac1. Note that Mac1 dataset comprises only actives.

